# A flexible photoacoustic retinal prosthesis

**DOI:** 10.1101/2024.09.03.611068

**Authors:** Audrey Leong, Yueming Li, Thijs R. Ruikes, Julien Voillot, Yuhao Yuan, Guo Chen, Clémence Bradic, Arnaud Facon, Chakrya-Anna Chhuon, Corentin Joffrois, Gilles Tessier, Marion Cornebois, Julie Dégardin, Jean-Damien Louise, Ji-Xin Cheng, Chen Yang, Hélène Moulet, Serge Picaud

## Abstract

Retinal degenerative diseases of photoreceptors are a leading cause of blindness with no effective treatment. Retinal prostheses aim to restore sight by stimulating residual retinal cells. Here, we present a photoacoustic retinal stimulation technology. We designed a polydimethylsiloxane and carbon-based flexible film that converts near-infrared laser pulses into a localized acoustic field with 56-µm lateral resolution, allowing precise stimulation of mechanosensitive retinal cells. This photoacoustic stimulation robustly and locally modulated retinal ganglion cell activity in both wild-type and degenerated *ex vivo* rat retinae. In animals subretinally implanted with a millimeter-sized photoacoustic film, pulsed laser stimulation generated neural modulation along the visual pathway to the superior colliculus, as measured by functional ultrasound imaging. The biosafety of the film was confirmed by the absence of short-term adverse effects , while local thermal increases were measured below 1 °C. These findings demonstrate the potential of photoacoustic stimulation for high-acuity visual restoration in blind patients.

## Introduction

Retinitis pigmentosa and age-related macular degeneration affect millions of people worldwide^1^, resulting in irreversible photoreceptor degeneration and blindness. Currently, there is no effective drug treatment for preventing photoreceptor loss. Retinal prostheses are implantable devices designed to stimulate the residual retinal layers to restore vision^2^. To date, two retinal prosthesis designs, which both primarily use electrostimulation to restore vision, have been approved for commercial implantation^3^. However, both were later removed from the market, facing challenges such as poor spatial resolution and small restored visual field. The more recent photovoltaic-based PRIMA implant, one of the most advanced retinal prostheses currently in clinical trials (NCT04676854), offers a spatial resolution of 100 µm, and a median visual acuity below the threshold for legal blindness^4^, and a limited restored visual field of 7°.

Focused ultrasound is a promising noninvasive approach for retinal stimulation^5^. It has been shown to evoke stable responses in the *ex vivo* salamander retina with a 90-µm lateral resolution at 43 MHz^6^ and in the *in vivo* blind RCS rat retina with a lateral resolution of around 400 µm with a 4.5-MHz transducer 2D-array^7–9^. However, it still faces safety challenges to meet FDA safety threshold for ophthalmological use^7^.

Photoacoustic modulation is an emerging method for high-precision neural stimulation^10,11^. A localized ultrasound field is generated by the photoacoustic agents upon excitation with a pulsed laser, and used to stimulate neurons. High spatial resolution stimulation of single neurons has been demonstrated through a fiber-based photoacoustic emitter^10^.

In this study, we investigated photoacoustic retinal stimulation as an alternative strategy for restoring vision. We here provide evidence of its efficacy both *ex vivo* and *in vivo* on healthy and degenerated retinae, as well evidence of the biosafety of the photoacoustic film in short-term implantation.

## Results

### Fabrication and characterization of the flexible photoacoustic film

The working principle of photoacoustic retinal stimulation is illustrated in Fig. 1a. A 1030-nm pulsed laser is delivered to the back of the eye onto the retina, illuminating the subretinally-implanted flexible photoacoustic (PA) film. Laser absorption by the film produces transient heating, which causes thermal expansion and compression of the material, thereby generating pulsed ultrasound waves. The generated ultrasound waves stimulate the mechanosensitive retinal cells^6,8,12^, resulting in a change in retinal ganglion cell activity, which is then transferred to the brain.

**Figure 1.**
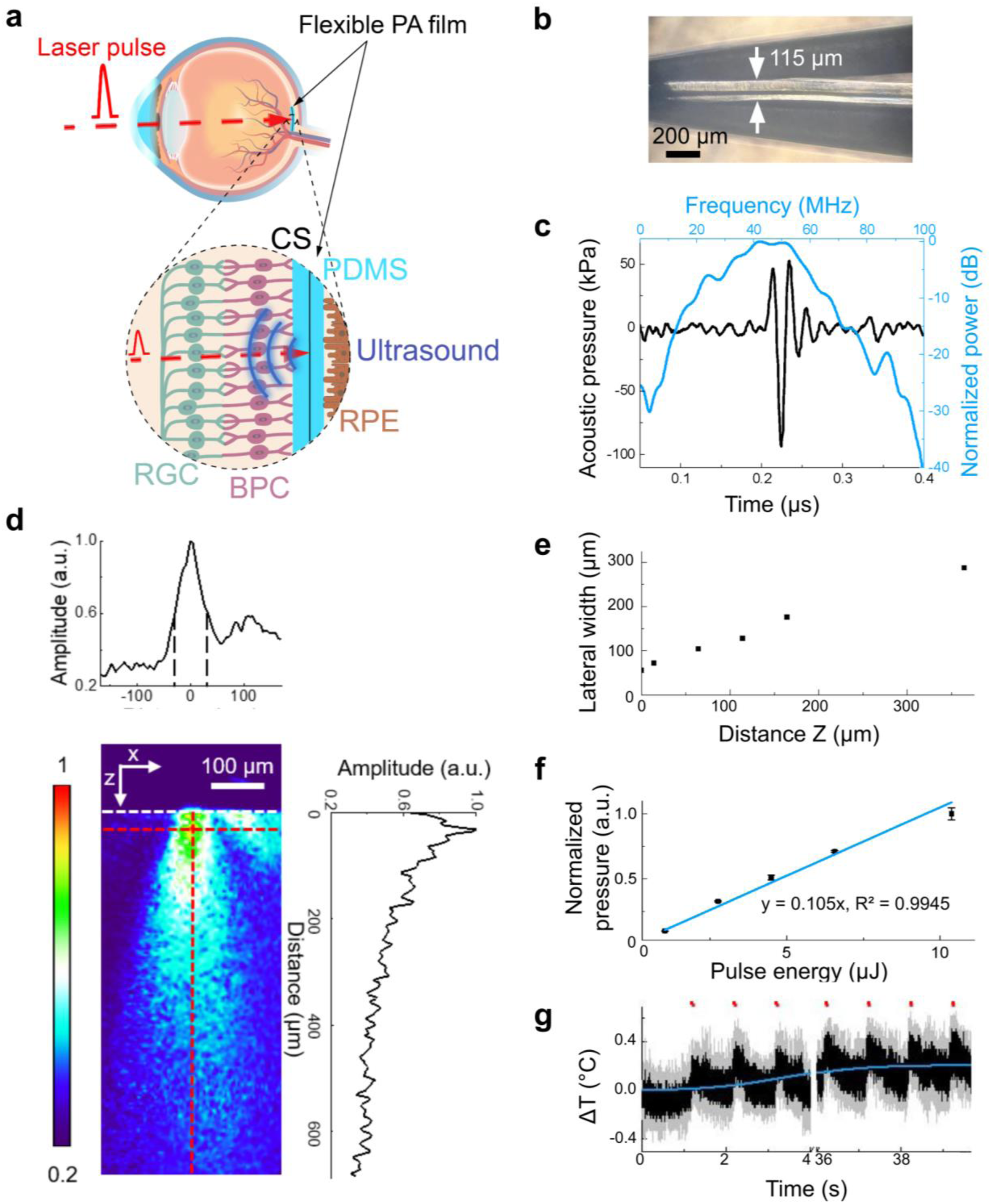
Characterization of the flexible photoacoustic film. **(a)** Working principle of the flexible photoacoustic (PA) film. Illumination of the PA film (cyan) with a nanosecond pulsed laser (red dashed line) produces an ultrasound emission (blue). CS: candle soot, PDMS: polydimethylsiloxane, RPE: retinal pigment epithelium, RGC: retinal ganglion cells, BPC: bipolar cells. **(b)** A photograph of the PDMS/CS/PDMS film with a three-layer design held by a tweezer. **(c)** Characterization of the PA film in the temporal domain (black) and frequency domain (blue) measured 0.9 mm away from the film surface. **(d)** Mapping of the ultrasound field generated by the PDMS/CS/PDMS film upon illumination through a 50-µm optical fiber. *Center*: measured distribution of the generated US field. The side lobe on the right of the field is due to the slight tilted angle when the optical fiber was put in contact with the sample film. White dotted line: interface between water and the film. *Top and right*: normalized lateral and axial profiles of the PA field, respectively, measured along the red dashed lines in the center panel. The amplitude of the acoustic signal was normalized to the maximum amplitude measured in the field. **(e)** Full width at half maximum of the lateral profile as a function of the axial position *Z* extracted from (d). **(f)** Peak-to-peak pressure of the PA signal as a function of laser energy per pulse measured from a PDMS/CS/PDMS film by a hydrophone. The pressure was normalized to the maximum pressure in all measurements. N = 3 for each data point. Blue line: linear fitting: y = 0.105x, R² = 0.9945. **(g)** Temperature increase at the surface of the PA film following illumination with a 200-µm laser spot. N = 3 for each data point, mean (black line) ± SD (gray shade). Red dots: laser on. Laser parameters: energy of 10 µJ per pulse, repetition rate of 3 kHz (laser power density P = 0.95 W/mm²), and burst duration 50 ms, delivered every 1 s over 40 s. The baseline change due to cumulative thermal effect was determined by logistic curve fitting of the data (blue line).

The PA film (Fig. 1b) consists of candle soot (CS) as the absorber material, sandwiched between two layers of polydimethylsiloxane (PDMS), which acts as the thermal expansion material. The film has a Young’s modulus of 2.12 ± 0.10 MPa to minimize the immune response^13,14^ once implanted (Supplementary Fig. S1). Upon excitation with 4.2-ns laser pulses at 7 µJ per pulse, the PDMS/CS/PDMS film emitted ultrasound pulses with a peak-to-peak pressure of 146.2 kPa measured 0.9 mm away (Fig. 1c). At the surface of the film, the conversion efficiency is estimated at 26 kPa.µJ^-1^ based on the distance-dependent pressure profile measured (Supplementary Fig. S2). PA signals were found to have a central frequency of 42.2 MHz and -6 dB bandwidth ranging from 29.6 to 59.9 MHz. This central frequency is an intrinsic property of our film under our experimental conditions, given the fixed laser pulse duration, the Grüneisen parameter of PDMS, and the constant concentration of the CS layer set by the constraints of the flame synthesis process. This central frequency has been demonstrated to activate e*x vivo* salamander retinae with a lower intensity threshold compared to lower acoustic frequencies^15^. These data suggest that the PDMS/CS/PDMS film is a promising photoacoustic converter for retinal stimulation.

The spatial distribution of the ultrasound field generated by the PDMS/CS/PDMS film was further mapped by PA-field microscopy (Fig. 1d, Supplementary Fig. S7). A 50-µm optical fiber was attached to the PA film to ensure a 50-µm-diameter illumination area. The axial pressure profile shows that the maximum acoustic signal is generated at the surface of the film upon illumination (*Z* = 0 µm) and attenuates to 50% of its peak value at *Z* = 140 µm (Fig. 1d, right). The lateral width (W) of the acoustic field, quantified by the full width at half maximum, measures W = 56 µm at Z = 0 µm and increases with axial depth to W = 124 µm at Z = 100 µm (Fig. 1e). These results confirm that under a confined illumination, the PA film produces a highly localized, 56-µm lateral ultrasound field comparable to the size of illumination, opening up potential for retinal stimulation with sub-100-µm resolution.

Measured acoustic pressure exhibited a linear relation with the incident laser energy per pulse (Fig. 1f), which indicates that the output pressure can be precisely modulated by adjusting the input laser energy.

The greatest heat increase is assumed to be generated at the level of the PA film, as a consequence of the laser light absorption. Ultrasound absorption could cause an additional increase in temperature, but it is expected to be much lower due to the very low acoustic energy. To ensure that the laser light absorption by the designed PA film is not associated with a substantial and detrimental temperature increase, we measured the temperature at the surface of the PA film. The tested laser conditions were consistent with those employed in the following *ex vivo* retinal stimulation experiments (next section). We observed a maximum temperature rise of 0.52 ± 0.09 °C (Fig. 1g). The baseline change due to cumulative thermal effects was 0.21 °C after 40 s. This value is an order of magnitude below the temperature increase needed for thermal neural modulation^16,17^ or tissue overheating. Therefore, the film is unlikely to thermally modulate retinal neuronal activity.

### Photoacoustic modulation of the *ex vivo* retina

To evaluate retinal responses following photoacoustic stimulation, we recorded the activity of retinal ganglion cells (RGCs) from *ex vivo* retinae of wild-type Long-Evans (LE) rats on a multi-electrode array (n = 4 rats). The PDMS/CS/PDMS film was placed on the *ex vivo* retina against the photoreceptor layer. It was photoactivated by a 1030-nm pulsed laser delivered through a 200-µm optical fiber (Fig. 2a), which was successively moved at different positions between stimulations. We applied 4.2-ns laser pulses at a repetition rate of 1.9 kHz (every 520 ns) for a burst duration of d_b_ = 10 ms, with a pulse energy of 10 µJ (Fig. 2b, top; power density P = 0.27 W.mm^-^²), yielding an estimated peak-to-peak ultrasound pressure of 0.12 MPa.

**Figure 2.**
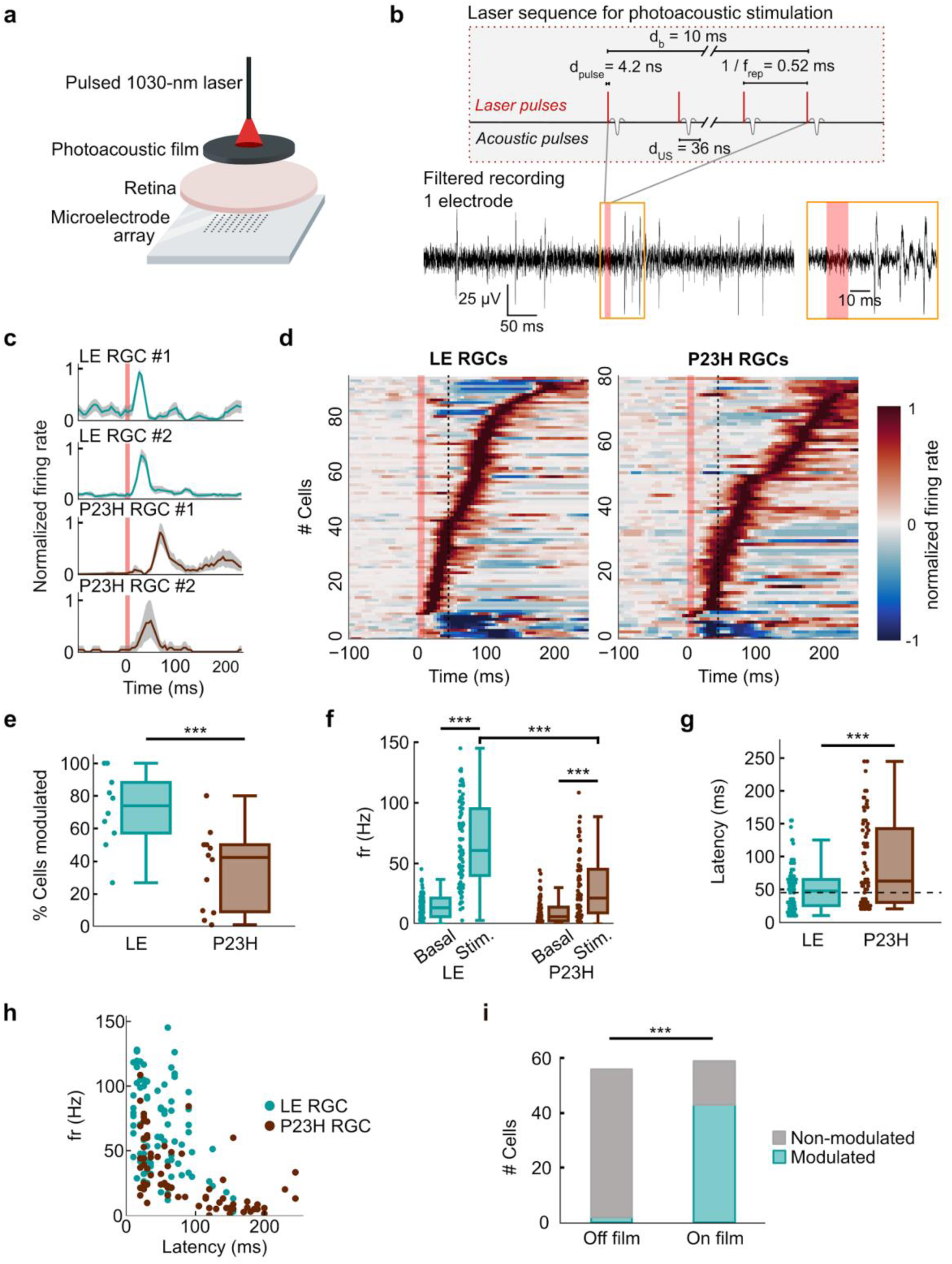
Photoacoustic modulation of *ex vivo* wild type and degenerated retinae. **(a)** The *ex vivo* retina was placed on a multi-electrode array (MEA) with the photoacoustic film against the photoreceptor layer. **(b)** *Top*: schematic of the laser sequence for photoacoustic stimulation. Laser pulses, with an energy of E_p_ = 10 µJ per pulse and duration d_pulse_ = 4.2 ns, were delivered at a repetition frequency f_rep_ = 1.9 kHz during a single burst of duration d_b_ = 10 ms. Each laser pulse is converted by the PA film into an acoustic wave with a duration d_US_ = 36 ns. *Bottom*: Example high-pass filtered MEA recording from a single electrode displaying elicited spikes following a photoacoustic stimulation. Red shaded area: laser on. Inset: zoom in of action potentials following stimulation. **(c)** Examples of Long Evans (LE) and P23H RGC responses to photoacoustic stimulation. Cyan (LE) and brown (P23H) lines: mean firing rate. Gray shaded areas: 99% bootstrapped CI from 1000 samples. Red shaded area: PA stimulation. **(d)** Heatmaps of normalized firing rates for responsive LE RGCs (left, n = 100) and P23H RGCs (right, n = 88). Red shaded area: PA stimulation. Dashed black line: 45-ms cutoff for slow and fast latency responses. Cells with excited responses display an increase in firing rate after photoacoustic stimulation (red), and cells with inhibited responses display a decrease in firing rate (blue). **(e)** Percentage of cells modulated by photoacoustic stimulation per stimulation site. LE: 74% (4 rats, n = 10 stimulation sites), P23H: 39 %, (4 rats, n = 12 stimulation sites). *** p < 0.001, Mann Whitney U test. **(f)** Firing rates of LE and P23H RGCs during baseline (basal) and following stimulation (stim). Mean firing rates: LE: fr_basal_ = 14 ± 1.0 Hz, fr_stim_ = 66 ± 3.7 Hz (n = 82 RGCs within stimulation range, p < 0.001, Wilcoxon signed-rank); P23H: fr_basal_ = 10 ± 1.2 Hz, fr_stim_ = 29 ± 2.88 Hz (n = 72 RGCs within stimulation range, p < 0.001, Wilcoxon signed-rank). **(g)** Latencies of RGC responses for LE (51 ± 34.2 ms, mean ± standard deviation) and P23H (89 ± 65 ms, p < 0.001 Wilcoxon rank-sum)). Dashed black line: 45-ms cutoff for slow- and fast-latency responses. **(h)** Firing rate of modulated RGCs as a function of response latency. Firing rate and response latency of excited RGCs were correlated for LE (r = -0.40, p < 0.001, Pearson correlation) and P23H (r = -0.59, p < 0.001, Pearson correlation) RGCs. **(i)** Off-film laser light stimulation. Percentage of LE RGCs modulated by direct laser stimulation on the retina (“off film”) compared to photoacoustic stimulation (“on film”). Laser parameters: E_p_ = 10 µJ per pulse, f_rep_ = 3.5 kHz, d_b_ = 10 ms. Off film: 3.6% ± 0.9% (n = 56 cells, 2 retinae), on film: 77% ± 20% (n = 59 cells, 3 retinae). Statistics: * p< 0.05, *** p < 0.001, Mann-Whitney U test.

Photoacoustic stimulation evoked robust RGC responses in healthy LE retinae (Fig. 2b, 2c, top two panels). Individual RGCs were considered responsive or modulated if their firing rate significantly increased (excited response) or decreased (inhibited response) relative to baseline (Fig. 2c, 2e). 100 LE RGCs (78%) exhibited an alteration in activity upon PA stimulation (Fig. 2e) out of the 129 spontaneously active RGCs on the electrodes contained in a 300-µm-radius area centered on the laser spot. Responsive RGC activity was mainly increased (92% of responding RGCs, Fig. 2d, left). RGCs with an increased activity had a mean response firing rate of 66 ± 3.7 Hz (Fig. 2f), and had a mean response latency of 51 ± 34.2 ms (Fig. 2g). The response latency was inversely correlated with the firing rate (Fig. 2h).

To investigate the potential of such photoacoustic stimulation for restoring vision, we then stimulated *ex vivo* retinae from blind P23H rats (n = 4 rats). Similarly to LE retinae, though with a lower fraction, 89 out of 229 P23H RGCs (39%) exhibited robust responses to photoacoustic stimulation (Fig. 2c-e), predominantly with increased activity (93% of responding RGCs). Compared to LE RGCs, P23H RGC firing rate was significantly lower following stimulation (29 ± 2.88 Hz, Fig. 2f) and response latency was significantly increased (89 ± 65 ms, Fig. 2g). Only 36% of P23H RGCs had response latencies below 45 ms^12^, which was significantly fewer than LE cells (39%, Fig. 2g). These results on the P23H rat retina demonstrate the *ex vivo* efficacy of photoacoustic stimulation in modulating retinal ganglion cells activity in degenerated retinae.

To rule out the possibility that the healthy LE retina was responding to the pulsed infrared laser light^18,19^, we applied direct laser pulses to the LE rat retina. Using identical laser conditions to those applied on the film, only 3.6 ± 0.9% of recorded RGCs showed a modified activity following directfilm stimulation, compared to 77 ± 20% of recorded RGCs following on-film photoacoustic stimulation (Fig. 2i). These results were further confirmed in *in vivo* LE rats (see below). Furthermore, we found that the PA film absorbs more than 99% of the infrared laser energy (Supplementary Fig. S3). These results confirmed that the infrared laser light filtered out through the implant was not sufficient to directly activate the retinal circuit in the healthy LE retina.

### Mechanistic exploration of retinal mechanosensitivity

It is currently unclear which layers in the retina contribute to the mechanosensitivity of the retina. We first observed an increase in response latency when photoreceptors have degenerated in P23H rat retinae, compared to LE retinae (Fig. 2g). To assess if photoreceptors initiate the short-latency US response, we bath-applied the group III mGluR agonist L-AP4 to LE retinae (Fig. 3a-c), which blocks the synaptic transmission between photoreceptors and ON bipolar cells^43^. We subsequently applied the kainate agonist ACET, which blocks most of the remaining glutamatergic synapses.

**Figure 3.**
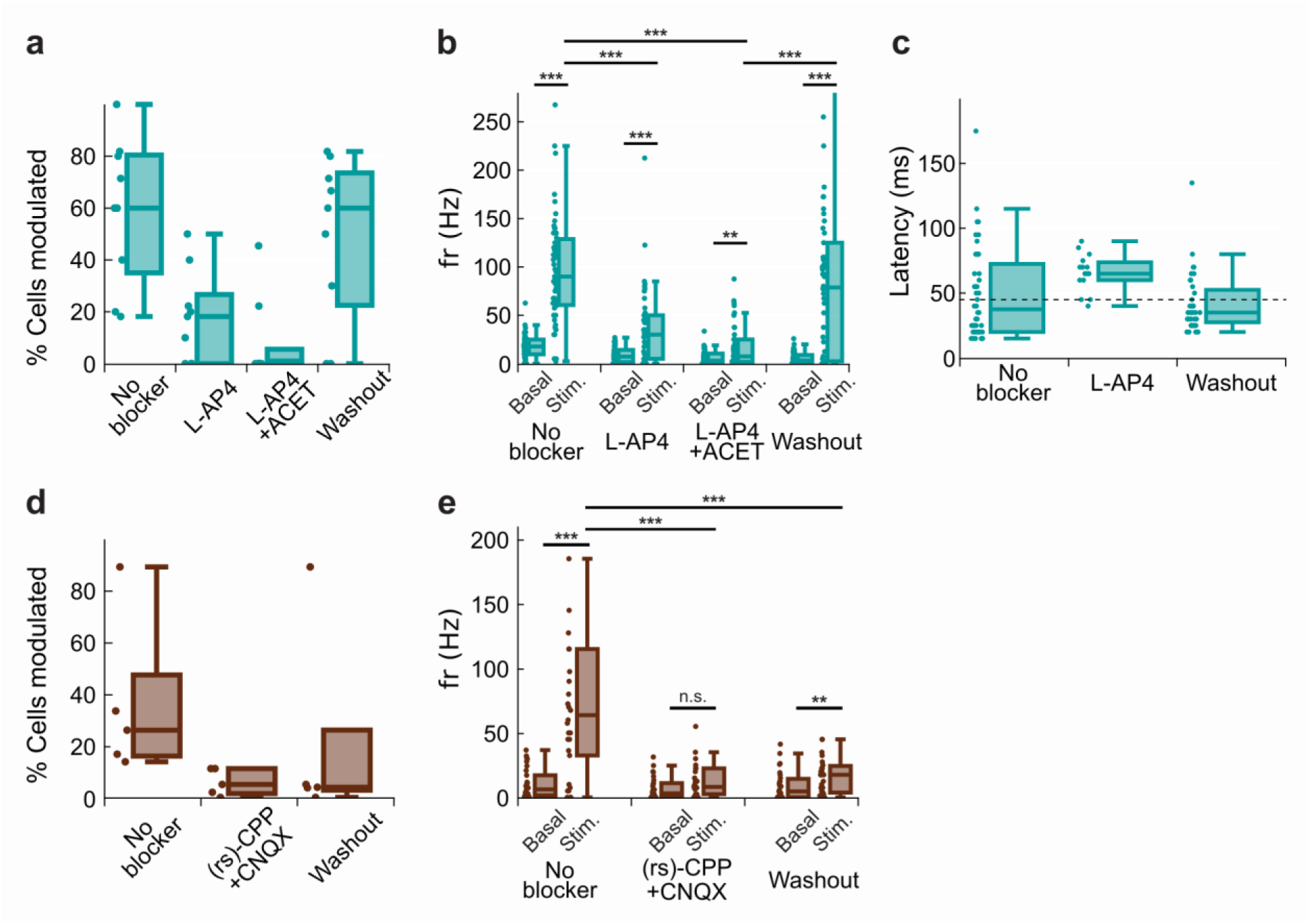
Photoacoustic responses under pharmacological blocking. **(a)** Percentage of cells modulated by photoacoustic stimulation per stimulation site before and after L-AP4 application. *No blocker*: 49%; *L-AP4*: 16%; *L-AP4 + ACET*: 7%; *washout:* 40% (2 retinae, n = 9 stimulation sites). **(b)** Group III mGluR agonist L-AP4 decreased RGC spontaneous firing rate and firing rate during PA-elicited responses in LE retinae. The population firing rate of RGCs (n = 41 cells, 2 retinae) is compared between baseline (Basal) and stimulation (Stim.), before blocker application (No blocker), following blocker application (L-AP4) and after washout (Washout) with Ringer medium. Firing rates per condition, *no blockers*: fr_basal_ = 18 ± 2Hz, fr_stim_ = 103 ± 9 Hz (p < 0.001); *L-AP4*: fr_basal_ = 9 ± 1 Hz, fr_stim_ = 37 ± 6 Hz (p < 0.001); *L-AP4+ACET*: fr_basal_ = 6 ± 1 Hz, fr_stim_ = 16 ± 3 Hz (p = 0.024), *washout*: fr_basal_ = 7 ± 1 Hz, fr_stim_ = 86 ± 13 Hz (p < 0.001). Comparison of the population stimulation firing rate fr_stim_ after blocker application and fr_stim_ before application (p < 0.001), and fr_stim_ after washout (p < 0.001). **(c)** L-AP4 increases response latency in LE retinae. Response latency per condition, *no blockers*: 49 ± 5 ms; *L-AP4*: 65 ± 4 ms; *washout*: 42 ± 3 ms. Comparison of the response latency before blocker application and after (p < 0.001, Wilcoxon signed-rank), and after washout (Friedmanchi square, p = 0.4). Due to absence of responses during application of L-AP4+ACET blockers, no latencies are reported for this group. Dashed black line: 45-ms cutoff for slow- and fast-latency responses. **(d)** Percentage of cells modulated by photoacoustic stimulation per stimulation site before and after (rs)-CPP and CNQX application in P23H retinae. *No blocker*: 42 %; *(rs)-CPP + CNQX*: 5 %; *washout:* 8 % (2 retinae, n = 5 stimulation sites). **(e)** Glutamate blockers (rs)-CPP and CNQX abolished RGC responses to PA stimulation in P23H retinae. The population firing rate of RGCs (n = 25 cells, 2 retinae) is compared between baseline (basal) and stimulation (stim), before blocker application (no blocker), following blocker application ((rs)-CPP+CNQX) and after washout (washout) with Ringer medium. Firing rates per condition, *no blockers*: fr_basal_ = 11 ± 2 Hz, fr_stim_ = 100 ± 23 Hz (p < 0.001); (rs)-*CPP+CNQX*: fr_basal_ = 7 ± 2 Hz, fr_Stim_ = 23 ± 10 Hz (p =0.141); *washout*: fr_basal_ = 8 ± 2 Hz, fr_stim_ = 24 ± 9 Hz (p = 0.016). Comparison of the population stimulation firing rate fr_stim_ after blocker application and fr_stim_ before application (p < 0.001, Wilcoxon signed-rank), and fr_stim_ after washout (p < 0.001, Wilcoxon signed-rank). For each panel, groups are tested for significant differences using Friedman chi square, with post-hoc Wilcoxon signed-rank. p-values are holm-corrected (***: p<0.001, **: p<0.05).

Before application of the blockers, a large fraction of recorded RGCs was responsive to the PA stimulation (58%, Fig. 3a, No blocker) with strong response firing rates (103 ± 2 Hz, Fig. 3a, No blocker). Following L-AP4 application, fewer RGCs were responsive to PA stimulation (19%, Fig. 3b, L-AP4), and their firing rates following stimulation were highly decreased (37 ± 6 Hz, Fig. 3b, L-AP4). Moreover, L-AP4 predominantly suppressed short-latency responses (<45 ms, 7% of recorded cells, Fig. 3c, L-AP4). These short-latency responses were recovered after the washout of the L-AP4 (Fig. 3c, baseline and washout): The L-AP4 condition mimics what was observed with recordings of P23H rat retinae with degenerated photoreceptors (p = 0.993, 2-sided Mann Whitney-U). These results further support that photoreceptors generate most of the short-latency RGC responses to ultrasound.

Addition of ACET to L-AP4 further suppressed RGC responses to PA stimulation (L-AP4 + ACET: 8 %, L-AP4: 19%, control with no blockers; 55%, Fig. 3b). This additional effect of ACET indicated that cells in the inner retina, upstream of RGCs, are also contributing to the production of the US-mediated RGC activation. These pharmacological data indicate that part of the RGC responses to PA stimulation rely on upstream singaling pathways. Taken together, these results demonstrate the important contribution of photoreceptors in the short-latency response to ultrasound stimulation, as shown by previous studies^6,18^, and reliance of PA-mediated stimulation on upstream signaling.

Finally, to isolate the contribution of RGCs in PA-induced responses in the P23H model, we bath-applied glutamatergic blockers (rs)-CPP and CNQX to P23H retinae (Fig. 3d, e). PA-induced responses were nearly completely abolished, but not recovered following the washout of the blockers (basal: 24 %, CPP+CNQX: 8 %, washout: 8 %, Fig. 3e). The lack of recovery during washout could be attributed to the difficulty of removing CPP and CNQX. These results suggest that the main mechanosensitive cells are upstream of RGCs, consistent with previous studies^6,12,18,19^, and that glutamate neurotransmission is required to transfer the mechanosensitive signal to the RGCs.

### Dependence of RGC response on laser conditions

We further investigated RGC responses to photoacoustic stimulation using different laser repetition rates (10 µJ per pulse, f_rep1_ = 1.9 kHz with P_1_ = 0.27 mW.mm^-^² and f_rep2_ = 3.5 kHz with P_2_ = 0.52 mW.mm^-^²) and burst durations (d_b_ = 5 - 30 ms). In LE retinae, the firing rates of cells with excited responses increased with burst durations up to d_b_ = 25 ms for f_rep1_, while at f_rep2_ they plateaued up to d_b_ = 15 ms and decreased with longer burst durations (Fig. 4a-b, left). P23H RGCs with excited responses also showed an increase in firing rate with longer burst durations for f_rep1_ (Fig. 4a-b, right), while firing rate increased for burst durations up to d_b_ = 20 ms then decreased with longer burst durations for f_rep2_ (Fig. 4b, right). LE RGC firing rates were significantly higher than those for P23H RGCs, up to 2.8-fold during d_b_ = 25 ms and up to 4.7-fold during d_b_ = 20 ms with f_rep1_ and f_rep2_, respectively (Fig. 4b). These results suggest that the degenerated retina requires stronger photoacoustic stimulation, consistent with previous findings concluding on a higher acoustic stimulation threshold in degenerated retinae compared to wild-type retinae^8^. LE RGC response latencies did not increase with burst duration or repetition rate (Fig. 4c), nor did P23H RGC latencies.

**Figure 4.**
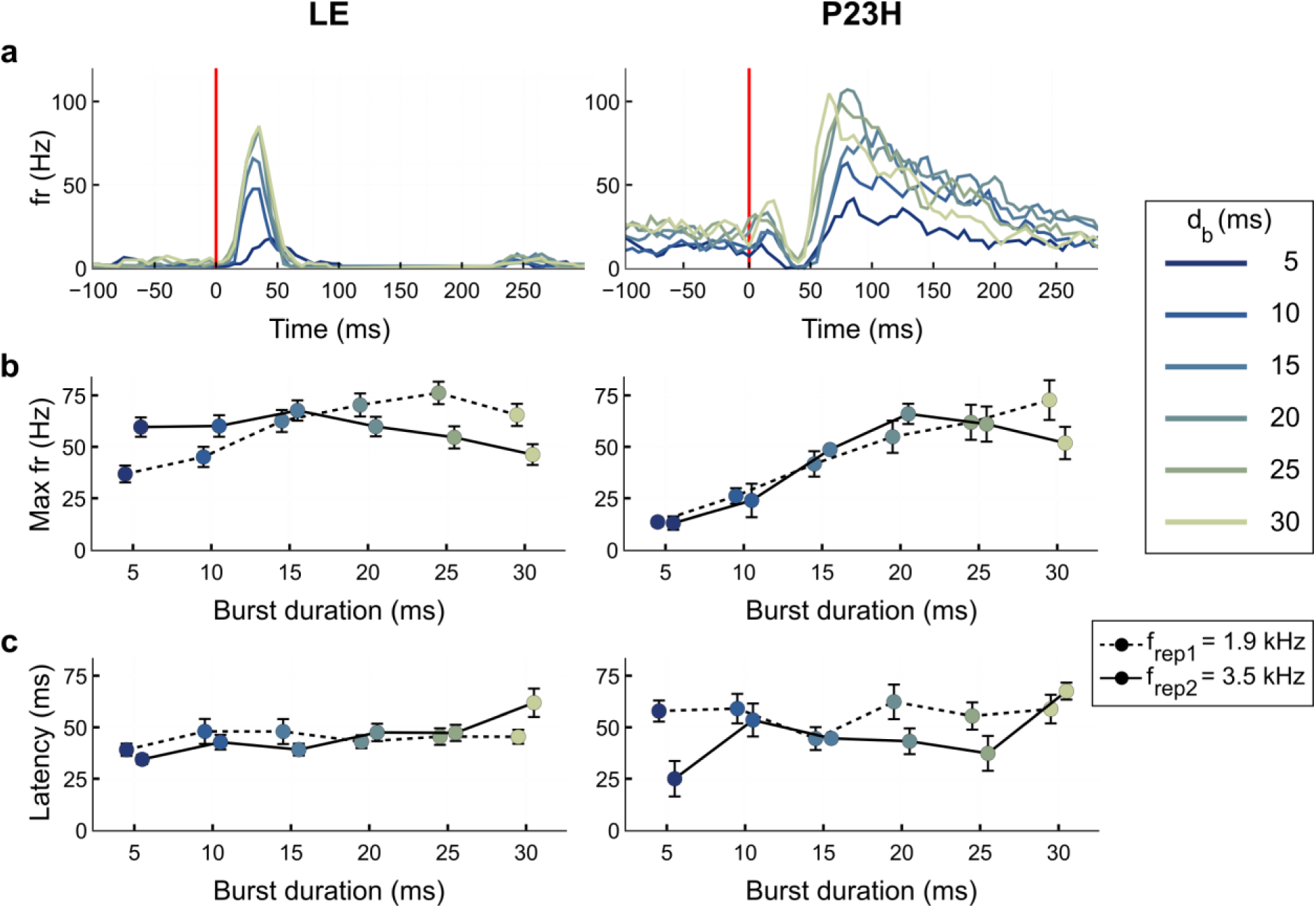
RGC responses under different laser burst durations and laser repetition rates. **(a)** Example LE (left) and P23H (right) cells showing increased maximum firing rate (fr) with increased burst duration (recorded at f_rep1_ = 1.9 kHz). Lighter colors indicate longer burst durations (d_b_= 5-30 ms). Vertical red lines: laser onset. **(b)** Maximum firing rate as a function of burst duration for LE and P23H RGCs during stimulation with repetition frequencies f_rep1_ = 1.9 kHz (dashed line) and f_rep2_ = 3.5 kHz (solid line). Data are plotted as mean + SE. In LE RGCs (left panel), the firing rate was positively correlated with burst duration for f_rep1_ (r = 0.91, p = 0.01, Pearson R). In P23H RGCs (right panel), the firing rate was positively correlated during both f_rep1_ (r = 0.996, p < 0.001) and f_rep2_ (r = 0.811, p = 0.05). With f_rep1_, for d_b_ = 5 ms and 20 ms, the maximum firing rate of LE RGCs was 2.8- and 1.3-fold higher, respectively, than for P23H RGCs (p < 0.001 for all conditions, Mann-Whitney U-test). With f_rep2_, for d_b_ = 5 ms, the maximum firing rate of LE RGCs is 4.7-fold higher than for P23H RGCs (p < 0.001, Mann-Whitney U-test). **(c)** Response latency as a function of burst duration for LE and P23H RGCs showed no significant correlation (LE: p = 0.70 and p = 0.19 for f_rep1_ and f_rep2_, respectively; P23H: p = 0.79 and p=0.61, Pearson R). In both (b) and (c), dashed lines: f_rep1_ = 1.9 kHz (P_1_ = 0.27 W.mm^-2^). Solid lines: f_rep2_ = 3.5 kHz (P_2_ = 0.52 W.mm^-2^). Dataset for (b) and (c): for LE, n = 244 cells, recorded from 4 retinae. For P23H, n = 104 cells, recorded from 4 retinae.

### Spatial resolution of *ex vivo* photoacoustic retinal stimulation

To investigate the spatial resolution of the photoacoustic retinal stimulation, we sequentially targeted multiple positions on the film by moving the laser-delivering 200-µm fiber at different sites above the photoacoustic film and mapped the responsive retinal cells (Fig. 5a). The laser repetition rate was set at f_rep1_ = 1.9 kHz for LE retinae (n = 11 sites) and f_rep2_ = 3.5 kHz for P23H retinae (n = 6 sites), to account for the higher modulation threshold previously described for P23H retinae.

**Figure 5.**
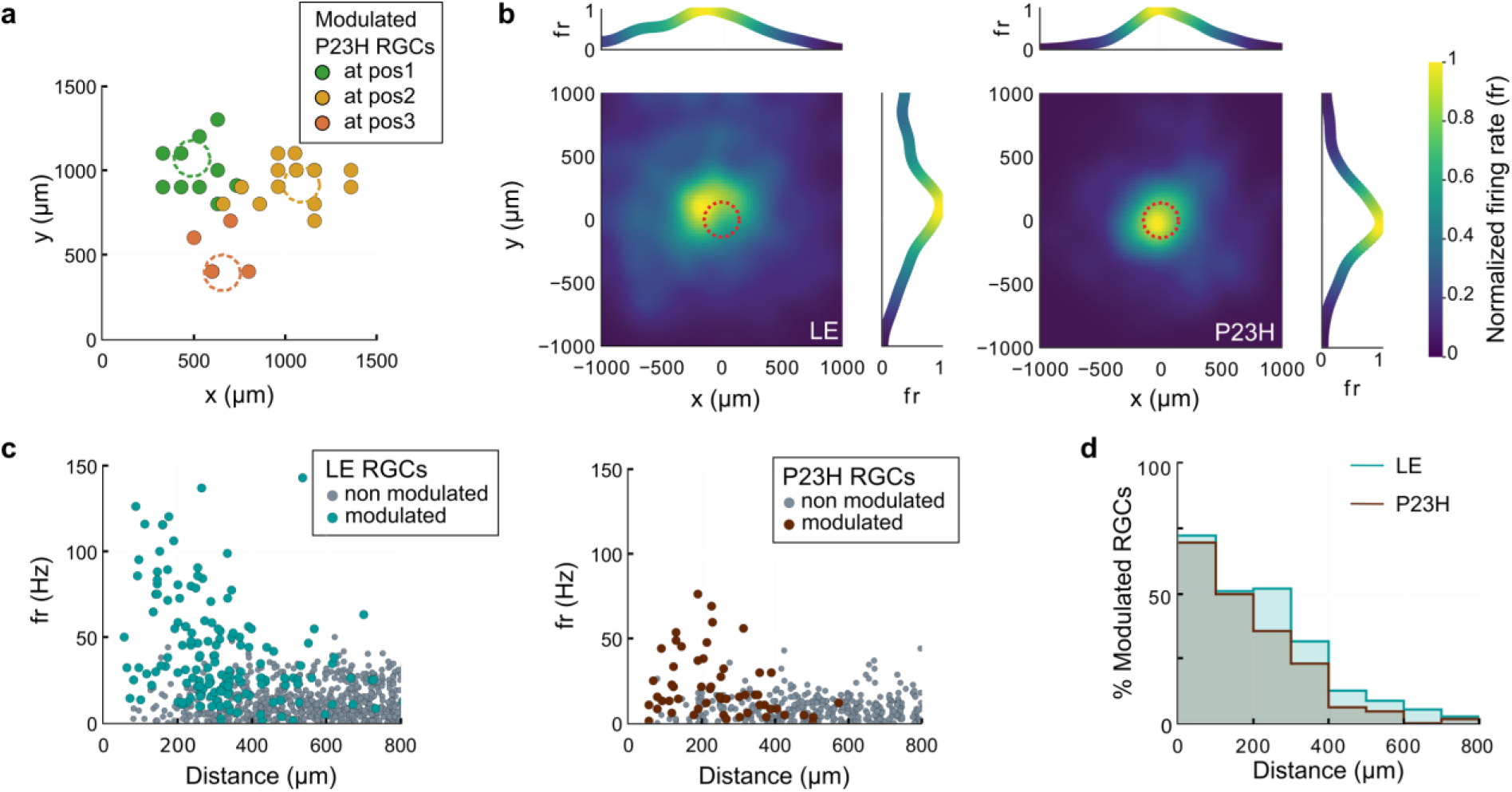
Spatial distribution of RGC modulation upon photoacoustic stimulation. **(a)** Different populations of RGCs were modulated by moving the laser fiber at different sites on the film. Example of a P23H retina stimulated at three sites; modulated cells at each stimulation site are grouped by color. The 300-µm-diameter laser spots are marked by dashed circles. **(b)** RGC firing rate of modulated cells normalized to maximum firing rate, mapped relative to the stimulation site for LE (left, 4 retinae, 11 stimulation sites) and P23H (right, 4 retinae, 6 stimulation sites). RGC maximum firing rates were averaged across all recorded cells at the same XY coordinates relative to the center of the laser spot (LE: n = 576 and P23H: n = 157 RGCs). Data were smoothed using convolution with a 100-µm gaussian kernel. Dashed circle: 300-µm-diameter laser spot. The shift between the maximum firing rate and the laser spot may be due to uncertainty in the laser spot coordinates, due to the 100-µm pitch of the MEA used for indirect measurement of the exact laser position. **(c)** Maximum firing rate for individual cells as a function of distance from the laser for LE (left) and P23H (right) RGCs. The response firing rate is negatively correlated with distance (LE: r = -0.310, p < 0.001. P23H: r = -0.268, p < 0.05, Pearson R). Each circle represents an individual cell. Cyan and brown: LE and P23H RGCs modulated by photoacoustic stimulation, respectively. Gray: non-modulated cells. **(d)** Percentage of RGCs modulated as a function of distance from the laser spot. Cyan: LE cells. Brown: P23H cells. Datasets for (c) and (d) are the same as for (b).

To assess the spatial distribution of PA-modulated RGCs with excited responses, we mapped the maximum RGC firing rate relative to the stimulation site (Fig. 5b). For both LE and P23H RGCs, the maximum firing rates were located within an area slightly larger than the laser spot (< 400 µm from center), and were negatively correlated with the distance from the laser spot (Fig. 5c). Furthermore, the percentage of responsive RGCs decreased when increasing the distance from the center of the laser spot (Fig. 5d). 73% of LE RGCs and 70% of P23H RGCs were modulated within a 100-µm distance, compared to 14% of LE RGCs and 6% of P23H RGCs at a 400-µm distance. These results indicate that stimulation with the PA film induces a localized response and demonstrate the potential for a high spatial resolution in photoacoustic stimulation.

### *In vivo* safety of the photoacoustic implants

To test the feasibility of photoacoustic stimulation and the biosafety of the film *in vivo*, we chronically implanted 1-mm-diameter PA films in the subretinal space of LE and P23H rats. To investigate whether there is a risk of increased degeneration of inner retinal layers with the 115-µm-thick PDMS/CS/PDMS film, we also prepared and studied a thinner PDMS-CNT film. The PDMS/CS/PDMS film, used in the previous *ex vivo* experiments, was optimized for high photoacoustic conversion efficiency and easy handling. A thinner 40-µm-thick uniformly mixed PDMS-CNT film (characterized in Supplementary Fig. S4) was designed to approach the 30-µm thickness of the clinically tested PRIMA photovoltaic implant, which has shown no long-term adverse effects aside from minor retinal thinning in patients^4^. Both types of films were treated with oxygen plasma to make them hydrophilic, in order to improve cell adhesion and thus minimize the risks of implant displacement.

After implantation, eye fundus imaging confirmed the correct positioning of the implant near the optic nerve and the overall integrity of the retina (Fig. 6a-b). No complications, such as retinal tearing after implantation or major inflammation after 7 days post-implantation (dpi), were observed on the OCT images and eye fundus exams. Although the presence of glial cells between the implant and the retina is expected, stimulation efficacy should not be affected, as acoustic attenuation is below 10 dB.mm^-1^ ^20^.

**Figure 6.**
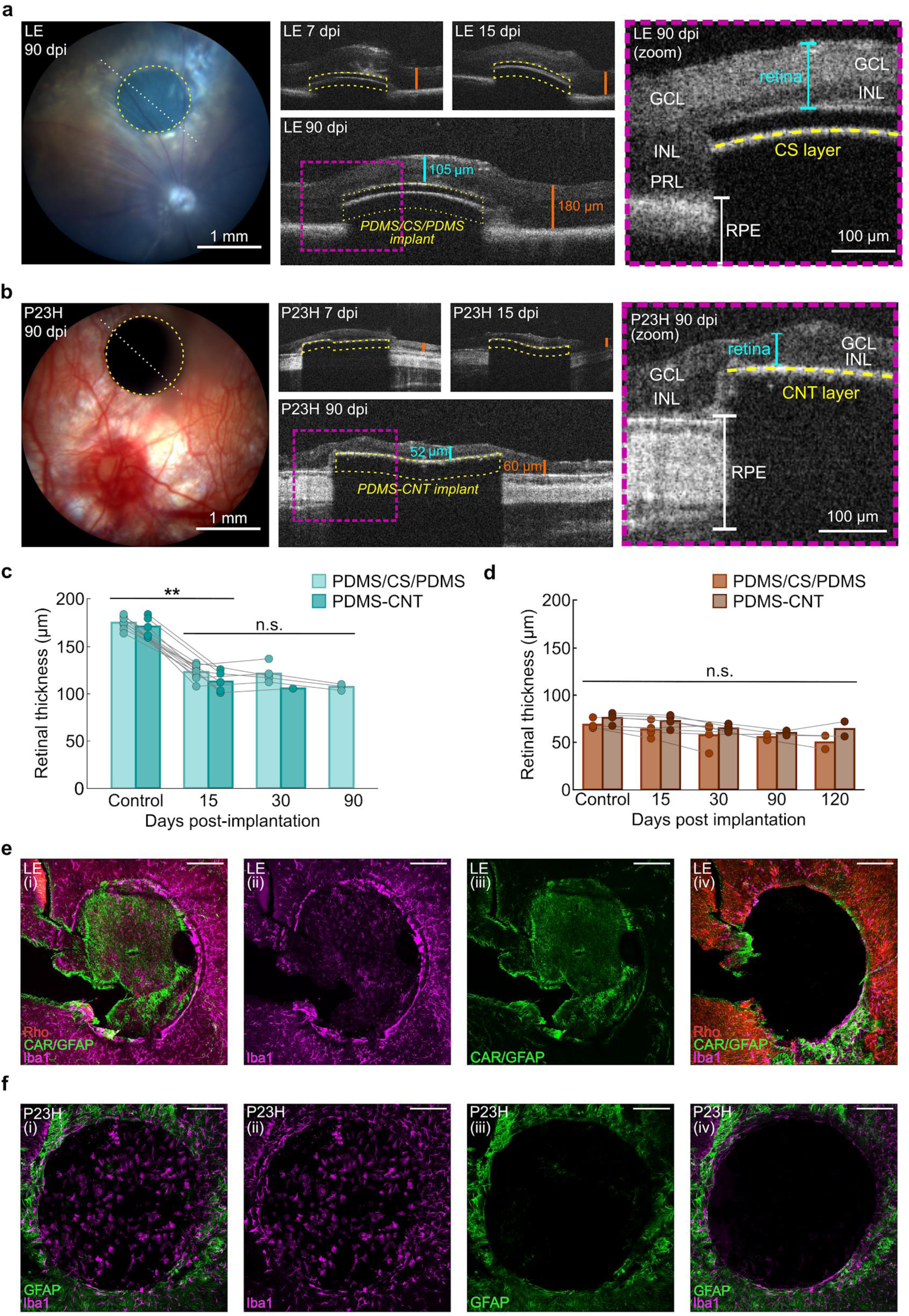
*In vivo* photoacoustic implant biocompatibility. **(a)** Eye fundus (left) and OCT images (middle and right) of an LE rat retina with a subretinal PDMS/CS/PDMS implant (yellow dotted line) at 7, 15, and 90 days post-implantation (dpi). OCT images were taken along the white dotted line shown in the left panel. In the zoomed-in OCT images, GCL: retinal ganglion cell layer, INL: inner nuclear layer, PRL: photoreceptor layer. RPE: retinal pigmented epithelium. The PR layer has degenerated above the implant. Right inset: zoom on the OCT image at 90 dpi. **(b)** Same as (a) but for a P23H rat. **(c)** Mean LE retinal thickness above PDMS/CS/PDMS (light cyan) and PDMS-CNT (dark cyan) implants over time. Control: mean retinal thickness next to the implant at 15 dpi. Thickness at 15 dpi and later is significantly lower than control thickness (**p < 0.01, Wilcoxon Signed-Rank test). At 15 dpi, the thickness above PDMS/CS/PDMS implants is not statistically different from the thickness above PDMS-CNT implants (p = 0.16, Mann-Whitney U test). Between 15 dpi and 90 dpi, the decrease in thickness above PDMS/CS/PDMS implant (123.2 ± 2.9 µm to 107.3 ± 2.0 µm) is not significant (p = 0.25, Wilcoxon Signed-Rank test). **(d)** Same as (c) for P23H rats. The difference of retinal thickness above both implants is not statistically significant (p = 0.16 at 15 dpi and 0.32 at 30 dpi, Mann-Whitney U-test). Retinal thickness is stable up to 120 dpi for both PDMS-CNT (p = 0.18, one-way ANOVA) and PDMS/CS/PDMS implants (p = 0.51). **(e)** Immunolabeling of an implanted LE retina, labeling rods (Rho, red, *i, iv*), cones (CAR, green, *i, iii*), microglia (Iba1, magenta, *i*, *ii*, *iv*), and Müller cells (GFAP, green, *i, iii*). *i - iii*: implanted zone. *iv:* zone surrounding the implant. **(f)** Same as (e) for P23H, with microglia (Iba1, magenta, *i*, *ii*, *iv*) and Müller cells (GFAP, green, *i, iii*). Scale bar for (e) and (f) is 200 µm.

On OCT, the average retinal thickness was 174.0 ± 2.3 µm for LE rats (n = 13) and 72.3 ± 2.3 µm for degenerated P23H rats (n = 8). At the PDMS/CS/PDMS implant position, LE retinal thickness decreased to 123.2 ± 2.9 µm at 15 dpi, 121.5 ± 4.2 µm at 30 dpi, and 107.3 ± 2.0 µm at 90 dpi (Fig. 6c). In LE rats implanted with the PDMS-CNT implant, similar values of 113.0 ± 4.8 µm at 15 dpi and 105.8 µm at 30 dpi were measured (Fig. 6c). The decrease in the retinal thickness above the implant in LE rats was likely due to photoreceptor degeneration (Fig. 6a, right), caused by the physical separation of photoreceptors from the retinal pigment epithelium, as previously reported with other prostheses^21,22^. In implanted P23H rats, retinal thickness above the implant remained stable and comparable to the neighboring area for up to four months for both PDMS/CS/PDMS and PDMS-CNT implants (Fig. 6d).

Finally, we performed immunohistochemistry assays on LE (n = 3) and P23H (n = 4) rats. We confirmed the absence of rods and cones at the level of the implant in LE rats (Fig. 6e, *i*). We also observed the presence of activated microglia at the implantation site in both LE (Fig. 6e, *ii*) and P23H rats (Fig. 6f, *ii*), and of activated Müller glia only in LE rats (Fig 6e, *iii*; P23H: Fig 6f, *iv*). No inflammation was observed in the zone surrounding the implant (Fig 6e-f, *iv*), except for a highly inflamed zone (Fig. 6e, *iv*, bottom right corner), which corresponded to the insertion track of the implant during surgery. Future studies will need to refine the implantation surgery of the PA implant to minimize inflammation caused by the insertion of the implant. Surface modification of the PDMS may also be explored to minimize inflammation at the level of the implant, in particular to reduce potential fibrosis.

### Photoacoustic retinal stimulation *in vivo*

We then examined *in vivo* photoacoustic stimulation of the degenerated retina using subretinal PDMS/CS/PDMS and PDMS-CNT implants in LE rats. As mentioned in the previous section (Fig. 6a, c), the local detachment of the retina from the retinal pigment epithelium (RPE) caused by the implant induces local degeneration of photoreceptors in LE rats at the location of the implant, creating a localized model of retinal degeneration at the photoacoustic stimulation site (Fig. 6e, f). Activation of the visual pathway was assessed in the contralateral Superior Colliculus (cSC) using functional ultrasound imaging (fUSI), which measures relative changes in cerebral blood volume (rCBV) triggered by neuronal excitation (Fig. 7a). We chose to record the cSC instead of the primary visual cortex (V1) since anesthesia adversely affects the capability of fUSI to detect activation in V1^30,31^. A cranial window was generated just prior to the experiment, preventing a long-term follow-up on the colliculus activity. To verify the coordinates of the cSC, we measured its natural visual activation using control full-field white light stimulation of the implanted eye (P = 0.02 mW.mm^-^²). This generated a large rCBV response area in the cSC (Fig. 7c, i), which was used as a reference area in further analyses (Fig. 7f). We measured a large increase in the amplitude of the rCBV, averaged on a 29-by-29-pixel (29 pixels = 302 ± 10 µm) square region of interest (ROI; Fig. 7f, light gray), which was centered on the maximum responses in the cSC. The same ROI was subsequently used to compute the averaged rCBV for all of the stimulation conditions (Fig. 7f). To better define the expected size of the activated area for PA stimulation, we focused a 400-µm-diameter spot of 595-nm laser light (P = 0.21 mW.mm^-^²) onto the healthy retina next to the implant (Fig. 7b, iii). It similarly triggered an increase in rCBV in the cSC (Fig. 7c, ii), with an activated area representing 32 ± 11 % (n = 6) of the area activated by the full-field white light (Fig. 7f). In the same 29-by-29-pixel ROI, the rCBV amplitude and response kinetics were similar to those generated by the full-field white light stimulation (Fig. 7g, orange).

**Figure 7.**
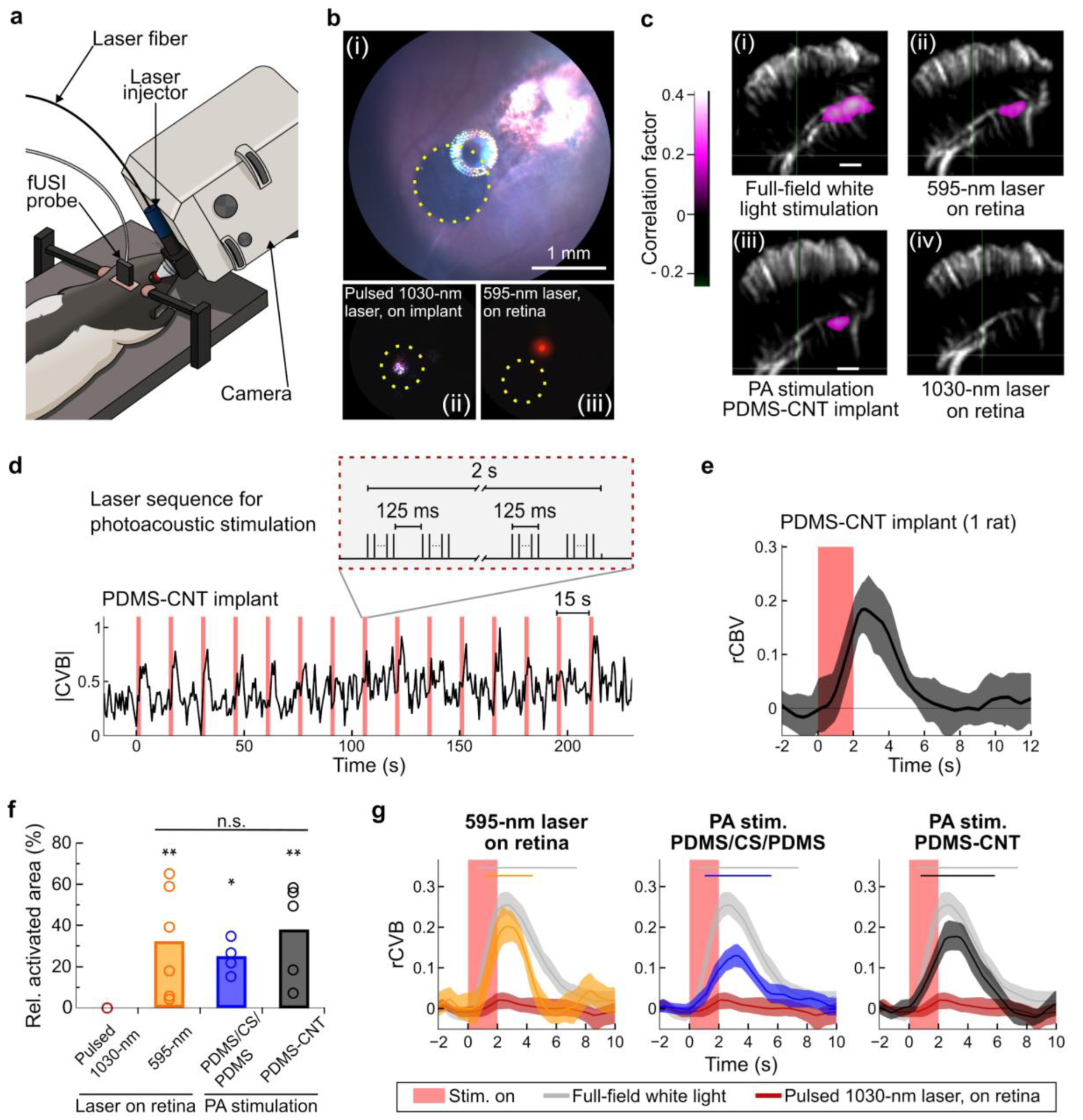
Superior colliculus activation following photoacoustic stimulation of *in vivo* LE retinae. **(a)** Setup for *in vivo* eye stimulation and fUSI recordings. **(b)** Eye fundus images of a 1-mm PA implant (*i,* yellow dotted circle) and 400-µm-diameter laser spots (*ii*: pulsed 1030-nm laser on the implant, *iii*: continuous 595-nm laser on the retina) used for laser and photoacoustic stimulation. **(c)** Functional ultrasound imaging in the coronal plane (left hemisphere, AP, -6.5 mm from the bregma). The correlation map displays the relationship between relative cerebral blood volume (rCBV) and the stimuli. Active pixels reflect regions of activated neurons in the contralateral superior colliculus (cSC) for a single recording (15 stimulations). **(d)** Top: Laser sequence for photoacoustic stimulation (repetition rate f_rep_ = 6.1 kHz). Bottom: example normalized CBV trace for pulsed 1030-nm photoacoustic stimulation on a PDMS-CNT implant (measured in 29-by-29-pixel region of interest of the cSC, chosen as the area with peak correlation to full-field-white-light stimulation). Baseline activity was recorded for 40 s before and after the stimulation session, and was used to normalize the CBV signal. **(e)** Average rCBV for a single session with 15 stimulations (same data as for (d)). Mean rCBV ± 99% CI. The laser sequence starts at 0 s. Red shaded area: PA stimulation. **(f)** Activated area following stimulation, relative to the area activated by full-field white light stimulation (rel. activated area), for different stimuli: full-field white light stimulation (light gray, 6 rats, n = 6 recordings), 1030-nm laser stimulation on the retina (dark red, 3 rats, n = 4 recordings), 595-nm laser stimulation on the retina (orange, 3 rats, n = 6 recordings), photoacoustic stimulation with PDMS/CS/PDMS implant (blue, 3 rats, n = 4 recordings), and photoacoustic stimulation with PDMS-CNT implant (black, 2 rats, n = 5 recordings). The activated area is measured by counting the number of pixels on correlation maps such as (c). Circles on the graph mark the ratio for individual recordings. Statistics: p-values vs white light stimulation: * p < 0.05, ** p < 0.01, Wilcoxon Signed-Rank test. PDMS/CS/PDMS vs 595 nm: p = 0.91, PDMS-CNT vs 595 nm: p = 0.792, PDMS/CS/PDMS vs PDMS-CNT: p = 0.56, Mann-Whitney U test. **(g)** Mean rCBV responses of individual rats following stimulation with a laser (595-nm and 1030-nm on retina) and photoacoustic stimulation, for all rats. Same data as (f). Horizontal bars denote significant elevation with respect to the baseline (e.g., no overlap of CI with basal CI). No significant difference in rCVB following photoacoustic stimulation between both implant types was found (e.g., overlapping confidence intervals). Peak rCBV values: white light, 0.26 (at 2.65 s); 595 nm, 0.20 (at 2.69 s); PDMS-CNT, 0.18 (at 3.24 s); PDMS/CS/PDMS, 0.13 (at 3.18 s); 1030-nm laser on retina, 0.02 (at 2.19 s). Shaded areas: 95% bootstrapped CI. Red shaded area: PA stimulation.

We then proceeded with photoacoustic stimulation on the implants (Fig. 7b, ii; Fig. 7d, top). The 1030-nm laser delivered eight 125-ms bursts during 2 s, repeated every 15 s, for a total of 15 stimulations per recording (Fig. 7d). Laser power densities were P = 0.29 ± 0.06 W.mm^-^² (mean ± SD) for PDMS/CS/PDMS implants and P = 0.39 ± 0.12 W.mm^-^² for PDMS-CNT implants. The estimated peak-to-peak acoustic pressures at the surface of the implants were 0.05 MPa and 0.15 MPa for PDMS/CS/PDMS and PDMS-CNT implants, respectively. We observed activation in a large cSC area following each photoacoustic stimulation (Fig. 7d-e) using both types of implants (Fig. 7c, iii). Photoacoustic stimulation with the PDMS/CS/PDMS and PDMS-CNT implants activated cSC areas measuring 25 ± 4 % (n = 4) and 38 ± 10 % (n = 5) of the full-field-white-light-activated area, respectively (Fig. 7f). Furthermore, in the 29-by-29-pixel ROI, the averaged rCBV amplitudes for the PDMS/CS/PDMS (Fig. 7g, blue) and PDMS-CNT (Fig. 7g, black) implant stimulations reached similar amplitudes to that generated with the 400-µm-diameter spot of the 565-nm laser light on the healthy retina (Fig. 7g, orange). The size of the activated areas generated by stimulation with the PDMS/CS/PDMS and PDMS-CNT implants, and with the 595-nm-laser spot, were not significantly different (Fig. 7f). By contrast, a direct 1030-nm-light stimulation of the healthy retina (Fig. 7c, iv; P = 0.56 ± 0.21 W.mm^-2^, 400-µm-diameter spot) did not generate a significant cSC activation (Fig. 7f). The averaged rCBV was negligible in the ROI, and statistically different from all of the other measurements (Fig. 7g, dark red). Taken together, these results show that the photoacoustic stimulation of the degenerated retina elicits a robust and local activation of the visual pathway downstream of the retina, with an amplitude and size comparable to a visible light stimulation of the retina.

## Discussion

In this study, we developed flexible photoacoustic films that efficiently generated acoustic waves. Although CS-PDMS films have been developed as laser ultrasound transducers^23–25^ and for superhydrophobic coating^26,27^, our study represents the first instance of engineering them into a sandwich-structured PDMS-CS-PDMS film as a biomedical implant, specifically designed to minimize heat deposition in the surrounding tissue and to minimize the diffusion of carbon particles into the biological tissue. The generated photoacoustic waves successfully stimulated the activity of retinal cells *ex vivo* and *in vivo*, thereby activating downstream visual pathways *in vivo*. This RGC stimulation was demonstrated both *ex vivo* and *in vivo* on degenerated retinae. These results therefore provide evidence that the degenerated retina maintains some ultrasound sensitivity, enabling RGC activity modulation and the consecutive stimulation of higher visual areas for visual restoration.

We showed that an acoustic pressure of 0.05 MPa was sufficient to elicit responses in the superior colliculus, which is two orders of magnitude lower than the pressure thresholds previously reported for *in vivo* transducer-based ultrasound retinal stimulation^9^. Pressure thresholds of photoacoustic stimulation correspond to mechanical indexes (MI) below 0.03 (PDMS/CS/PDMS film) and 0.1 (PDMS-CNT film) and spatial peak temporal average intensities (I_SPTA_) below 0.06 and 0.9 mW.cm^-^², respectively (Supplementary Table T1). Temperature increases were measured to be below 1 °C at the film surface (Fig. 1g, Supplementary Fig. S6). These metrics align with FDA safety guidelines for ultrasonic ophthalmic devices^28^, making our PA implants a promising option for safe ultrasound stimulation of the retina. Due to the bidirectional nature of our implant, acoustic waves are transmitted not only to the retinal side, but also to the opposite side toward the RPE and choroid, potentially impacting mechanosensitive elements therein. While the implications of ultrasound waves to the RPE and choroid require further investigation, the applied acoustic pressures are below safety thresholds and the acoustic waves are applied at 0.025% duty cycle. Thus limited or no aversive effects are to be expected on the RPE and choroid.

Restoring meaningful vision requires high spatial resolution over a large area of the retina. To achieve this, each individual stimulation source (pixel) must produce tightly confined fields with sub-20-µm^29^ lateral resolution, and maximize pixel density to 2500 px.mm^-^² in the macula, which covers a 25 mm² area in humans. Among current electrical retinal prostheses, complex 3D honeycomb photovoltaic devices^32^ and optimized laser stimulation sequences^33^ show potential to generate 20-µm-wide electric fields with dense arrays of thousands of electrodes, but each rigid implant can only cover at best 9 mm² of the retina. Ultrasound retinal stimulation offers the advantage of non-invasive stimulation over an area greater than the macula, but the pixel density of transducer 2D-arrays is very limited (currently 16 x 16^9^) regardless of the lateral resolution of the stimulation field, which remains above 80 µm (*in silico* data, at 20 MHz^9^). The photoacoustic film demonstrated in this study offers the potential to meet both of these needs due to its spatially continuous nature and the multiplexing capability of photons. We demonstrated through PA-field mapping that a 50-µm-diameter laser spot generates a 56-µm lateral ultrasound field, and that the RGC activity was locally modulated around the laser spot in both healthy and degenerated retinae. Neuronal spatial resolution based on MEA recordings still remains to be determined. Additionally, the potential of the photoacoustic film to enable high visual acuity using smaller laser spot sizes is currently under investigation. Moreover, the pixel number could exceed a million by using digital micromirror devices to project a laser pattern onto the continuous film. Whole-eye models will be necessary to predict the actual resolution that could be achieved for *in-human* applications. A comparison between our approach and other vision restoration technologies across multiple parameters is summarized in Supplementary Table T2.

Since the structures involved in retinal mechanosensitivity are not yet known, it is difficult to evaluate the potential adaptation or depression of mechanosensitivity to long-term ultrasonic retinal stimulation for vision restoration. It has been shown that retinal stretch with increased intraocular pressure (IOP) resulted in an increased expression of Piezo1 and 2 in RGCs^25^. While the similarities between the mechanisms underlying elevated IOP and ultrasonic mechanostimulation are not yet understood, it is conceivable that ultrasonic mechanostimulation may also lead to an increase in the expression of mechanosensitive proteins. In the course of our studies, no such depression was observed, but the stimulations were limited in time. Future studies need to be conducted to better evaluate the effect of long-term mechanical stimulation of the retina.

While further studies are required to investigate the mechanisms of photoacoustic retinal stimulation and whether the implant can restore meaningful vision to patients afflicted by retinal degenerative diseases, our results collectively demonstrate that photoacoustic retinal stimulation through flexible implants opens up potential for an innovative strategy for restoring vision, with high precision and a large field of view.

## Material and methods

### Design of the photoacoustic film

Both the laser wavelength and the PA materials were optimized for safety and performance. Light wavelengths ranging from 500 nm to 1150 nm have maximum transmission in the human eye media^34^. A nanosecond laser with a 1030-nm wavelength was chosen to maximize transmission to the retina^34^, while avoiding triggering responses in photoreceptors, as AMD patients may retain peripheral vision. CS and CNT were selected as the absorber material due their high photoacoustic conversion efficiency, accessibility, and lower safety concerns compared to lead-containing materials^35,36^. For the thermal expansion material, PDMS was identified as the best option due to its transparency, high Grüneisen parameter, excellent biocompatibility, and stability^37^. Although CS-PDMS films have been developed as laser ultrasound transducers^23–25^, our study represents the first instance of engineering them into a sandwich-structure. The PDMS mixing ratio was adjusted to 5:1 to increase the Young’s modulus, thereby enhancing photoacoustic conversion efficiency^38^. The central frequency of our film is an intrinsic property of our film under our given experimental conditions. Several factors influence the ultrasound frequency generated by a photoacoustic device, including the laser pulse duration, the Grüneisen parameter of the elastomer material, and the effective optical absorption depth. In our setup, the laser pulse duration is fixed, PDMS is the only choice of absorber because it is both transparent and exhibits a high Grüneisen parameter, and the concentration of the CS layer remains constant due to the constraints of the flame synthesis process.

### Fabrication of the photoacoustic films

To fabricate the 115-µm-thick PDMS/CS/PDMS film, a uniform layer of candle soot was flame-synthesized and deposited onto a glass slide for 20 s, achieving a thickness of approximately 4 µm. Subsequently, a degassed PDMS mixture of silicone elastomer base and curing agent (Sylgard 184, Dow Corning Corporation, USA) with mix ratios of 5:1 was spin-coated at 500 rpm onto the candle soot layer, and cured at 110 °C for 15 minutes. The resulting cured film was then detached from the glass slide, inverted, and reattached. Another layer of PDMS mixture was spin-coated at 500 rpm and cured at 110 °C for 15 minutes.

To fabricate the 40-µm-thick PDMS-CNT film, we employed a recipe derived from previous work^10^. We initially prepared PDMS at mix ratios of 10:1. Subsequently, a 15%wt of CNT (<8 nm OD, 2–5 nm ID, length 0.5–2 µm, VWR, Inc., USA) was mixed with the PDMS, with the addition of IPA to facilitate CNT dissolution. The resulting mixture underwent a 5-minute sonication process, followed by a 30-minute degassing step to eliminate bubbles and IPA. The prepared mixture was then spin-coated onto a glass substrate at 500 rpm for 5 minutes. The coated substrate was cured at 110 °C for 15 minutes.

Both PDMS/CS/PDMS and PDMS-CNT films were treated with oxygen plasma for 1 min on both sides, to make the implant surface hydrophilic^39^. This allows for better cell adhesion, which reduces the risks of film displacement after the implantation surgery. The film was cut into smaller areas (5 x 5 cm²) and stored in distilled water before use to avoid reversion to their hydrophobic state. Biopsy punches (Kaimedical) of 1 mm and 1.5 mm were used to cut individual photoacoustic films for *ex vivo* and *in vivo* experiments.

### Characterization of the photoacoustic properties of the films

The photoacoustic properties of the films were characterized with a 40-µm needle hydrophone system (NH0040, Precision Acoustics Inc., UK) or an 85-µm needle hydrophone system (HGL-0085, Onda Corporation, USA). Illumination was provided by a Q-switched diode-pumped laser with a pulse width of 8 ns (RPMC, wavelength 1030 nm, repetition frequency 2.9 kHz, USA), delivered to one side of the film via a multimode fiber with a 200-µm core (FT200UMT, Thorlabs, USA). On the other side of the film, the hydrophone was mounted on a 3D stage and aligned with the area illuminated by the optical fiber. The signals were amplified with a pulser-receiver (Olympus, Model 5073PR, USA) and then recorded via a digital oscilloscope (Rigol, DS4024, USA).

### Mapping the photoacoustic pressure field

Photoacoustic field microscopy was used to map the generated ultrasound field (Supplementary Fig. S7), as previously reported^40^. Here a 1064-nm pulsed laser (OPOLETTE 355 LD, OPOTEK, pulse duration 5 ns) was used as the pump beam. A continuous wave 1310-nm laser (1310LD-4-0-0, AeroDIODE Corporation) served as the probe. A piece of PDMS/CS/PDMS film was mounted on a 50-µm optical fiber (FG050UGA, Thorlabs), and the 1064-nm laser was delivered to the film sample to generate the photoacoustic signals. A translation stage (ProScan III, Prior) was used to scan the generated ultrasound field. Under a single ns pulse, the PA-induced refractive index change was detected as the imaging contrast.

### Temperature measurements

A J-type thermocouple with a 200-µm tip was set against the PDMS/CS/PDMS film inside 3% agarose gel, typically used for mimicking tissue^41^. The PDMS/CS/PDMS film was attached to a 200-µm optical fiber to assure the alignment between the illuminated area on the film and the thermocouple tip. A Q-switched diode-pumped laser with a pulse width of 4.5 ns (RPMC, wavelength 1030 nm, USA) was used to illuminate the film. The temperature on the film was recorded with a 2 kHz sampling rate from 10 recordings. Average data was computed from the 3 recordings that showed the highest temperature rise. Photos of the setup are shown in Supplementary Fig. S5. No transient temperature events faster than the 2 kHz acquisition frequency are expected to occur (Supplementary Fig. S6).

### Animals

All animal experiments were conducted at the Paris Vision Institute, in accordance with the National Institutes of Health Guide for the Care and Use of Laboratory animals. Protocols were approved by the Local Animal Ethics Committee (Committee Charles Darwin CEEACD/N°5, project reference Apafis#40263-2023010909277429 v5) and conducted in agreement with Directive 2010/63/EU of the European Parliament. Wild-type Long-Evans male rats aged between 2 and 8 months were obtained from Janvier Laboratories. P23H male and female transgenic rats (9-14 months old) were raised locally. P23H rats serve as a model for autosomal dominant retinitis pigmentosa^42^.

### *Ex vivo* experiments

#### *Ex vivo* retina preparation and blockers

The following procedures were carried out under dim red light. Animals were dark adapted for 30 minutes, then anesthetized with CO_2_ and euthanized by cervical dislocation. The eyes were enucleated and hemisected in carboxygenated (95% O_2_, 5% CO_2_) Ringer medium containing (in mM): 125 NaCl, 2.5 KCl, 1 MgCl_2_, 1.25 NaH_2_PO_4_, 20 glucose, 26 NaHCO_3_, 1 CaCl_2_ and 0.5 L-Glutamine at pH 7.4. The medium was continuously perfused in the recording chamber at a speed of 1.5 mL.min^-1^ and was kept around 37 °C.

Isolated retinae were placed on a dialysis membrane (Spectra/Por® 6 50 kD dialysis membrane, Spectrum) coated with poly-L-lysine (0.1%, Sigma), with the photoacoustic film between the dialysis membrane and the retina, and with photoreceptors against the film. The retinae were pressed against a multi-electrode array (MEA) (MEA256 iR-ITO; Multi-Channel Systems, Reutlingen, Germany) with a custom 3D-printed piece.

Pharmacological blockers were bath-applied through the perfusion line. AMPA/kainate glutamate receptor antagonist 6-cyano-7-nitroquinoxaline-2,3-dione (CNQX, 20 μM, Tocris Bioscience) and NMDA glutamate receptor antagonist (RS)-3-(2-carboxypiperazin-4-yl)-propyl-1-phosphonic acid ((RS)-CPP, 10 µM, Tocris Bioscience) were used to block signaling upstream of RGCs and therefore isolate the contribution of RGCs in PA-induced responses. Group III metabotropic glutamate receptor agonist L-(+)-2-Amino-4-phosphonobutyric acid (L-AP4, 20 µM, Tocris Bioscience) was used to block the signaling between photoreceptors and ON bipolar cells^43^ to better understand the contribution of photoreceptors in retinal mechanosensitivity. Kainate antagonist (*S*)-1-(2-Amino-2-carboxyethyl)-3-(2-carboxy-5-phenylthiophene-3-yl-methyl)-5-methylpyrimidine-2,4-dione was used to block remaining PA responses after L-AP4 application.

#### *Ex vivo* photoacoustic retinal stimulation

Photoacoustic stimulations were done with a 1030-nm, 4.2-ns-pulsed laser (One DPSS, Bright Solutions) delivered through a 200-µm-core 0.22 NA multimode SMA/SMA optic fiber (Thorlabs Inc, USA., ref M25L01). A second 200-µm-core was connected to the first fiber using a fixed attenuator (Thorlabs Inc., USA, ref FA26M) to control the power density. The optical fiber was inserted into a custom 3D-printed holder incorporated in a motorized XYZ stage with 5-nm precision (Sensapex, uMp-3 micromanipulator). It was lowered above the PA film at a ∼90° angle and placed at a fixed (∼ 1 mm) distance above the PA film. The illumination spot was measured ∼300 µm in diameter on the film using ImageJ and the MEA electrode pitch as reference. A low power 650-nm guiding beam (FIBERCHECK, Laser Components) was used to calibrate the beam position relative to the MEA.

Laser pulse repetition rate and the laser burst trains were controlled using a Teensy microcontroller custom written software (C++, Java, Python). In a typical stimulation, the laser delivered 10 µJ pulses with a repetition frequency f_rep_ of 1.9 kHz or 3.5 kHz during a single 5-ms to 30-ms burst, repeated at 1 Hz for 40 bursts. PA film integrity was confirmed by the lack of photoelectric effect in the MEA recordings (Supplementary Fig. S8).

#### Analysis of MEA recordings

MEA raw traces were recorded through the MEA software (MC Rack, Multichannel Systems). Spikes were sorted using SpyKING CIRCUS^44^, manually curated using phy^45^, and attributed to individual cells. Spikes were referenced relative to stimulus onset and grouped across trials in bins using a sliding window (bin width = 20 ms, increments = 5 ms). Cell activity in each bin was estimated using bootstrap resampling (n = 1000 resamples, 99% confidence intervals), and considered significantly increased or decreased if there were no overlapping confidence intervals compared to baseline (200 - 100 ms before stimulus onset).

RGCs were considered responsive or modulated if their firing rate was significantly increased or decreased compared to baseline for at least 15 ms consecutively, and response latency was defined as the first bin of this series. Noise clusters were filtered from the cell clusters by excluding cells with response latencies below 5 ms. A 300-µm-diameter area was illuminated by the 1030-nm laser (200-µm fiber) during photoacoustic stimulation. For quantifying responsive cells and dose responses (Fig. 2 and 3), we included only cells within 300-µm of the center of the illuminated area (“stimulation site”). The number of recorded cells per stimulation site is shown in Supplementary Fig. S9. Cells with a response latency above 250 ms were excluded, as they were likely not a result from direct stimulation.

To analyze the relation between cell modulation and distance from the stimulation area (Fig. 5a), we calculated local averages in firing rate by assigning the firing rate of each cluster to a bin in a grid with a spacing of 25 µm, and smoothing the resulting averages using a convolution with a gaussian kernel (sigma = 100 µm).

#### *In vivo* experiments

Successful implantation was defined as good positioning of the 1-mm-diameter film in the subretinal space, with no complications due to surgery. N = 8 adult (9-10 mo) P23H rats were successfully implanted and used for biocompatibility studies. N = 7 adult Long-Evans rats were successfully implanted at 8 weeks of age and used for photoacoustic stimulation.

#### Surgery procedures for chronic subretinal implantation

A 1-mm-diameter PA film was surgically placed in the subretinal space in the central region next to the optic nerve, as previously described^31^. Briefly, a small sclerotomy was performed on the dorsal sclera tangential to the cornea. A gel of sodium chondroitin sulfate-sodium hyaluronate (Viscoat Alcon) was injected in the sclerotomy to generate a retinal detachment. The implant was then inserted below the detached retina in the subretinal space, targeting a location adjacent to the optic disk.

#### Ocular imaging

Eye fundus imaging (MICRON® IV, Phoenix, USA) and optical coherence tomography (Bioptigen® OCT system, Leica microsystems, Germany) were performed at 7 and 15 days post-implantation (dpi) for all rats to monitor inflammation and confirm correct implantation. Additional imaging was conducted at 30, 60, 90 and 120 dpi for rats that did not undergo prior retinal stimulation.

#### Cranial window acute surgery

Anesthesia was provided with an intraperitoneal injection of 40 mg.kg^-1^ ketamine (Axience, France) and 0.14 mg.kg^-1^ medetomidine (Domitor®, Vétoquinol, France) diluted in sodium chloride. The animal was placed on a stereotaxic frame to perform a left craniotomy. Drops of ocular gel (Lubrithal®, Dechra, France) were applied and the eyes were then covered with a black cloth for dark adaptation. A rectangular piece of bone was removed from Bregma -3 mm to -8 mm.

#### Retinal stimulation and brain imaging

Retinal stimulation was performed 26 - 40 days after implantation surgery for rats with PDMS/CS/PDMS implants and 23 - 29 days after implantation surgery for PDMS-CNT implants, and immediately after the cranial window surgery. Anesthesia was re-administered every 45 min with one-third of the initial dose, up to a maximum of 5 injections. At the end of the experiment, the animals were euthanized using an intracardiac injection (Exagon®, Axience, France).

For full field light stimulations with a white LED source, the light power on the retina was estimated to be ∼0.02 mW.mm^-^², based on the power entering the pupil of 1.2 mW. The choice of the stimulation protocol was informed by prior retina studies using functional ultrasound imaging^31,46^. Each 1.8-s stimulation sequence consisted of 6 evenly spaced 300-ms illuminations (LED on), repeated 15 times.

For 595-nm, 1030-nm light stimulation and photoacoustic stimulation, focused laser spots were aimed using a laser injector from the MICRON® 810-nm Image-Guided Laser modality combined with a MICRON® III camera (Phoenix, USA). A low power 650-nm guiding beam (FIBERCHECK, Laser Components) was coupled to the injector to safely choose the area to stimulate. The rat’s implanted eye was covered in ocular gel (Lubrithal®, Dechra, France) and in contact with the camera lens. Stimulation sequences for all 3 modalities were identical.

For 595-nm (continuous) laser stimulation, power density on retina was 26 µW in a 400 ± 26 -µm-diameter laser spot. For photoacoustic stimulation, the same 1030-nm pulsed laser used in the *ex vivo* experiments was employed. The laser energy exiting the laser injector was E_p_ = 15 µJ per pulse. To aim at the implant for photoacoustic stimulation, the laser focal spot was not placed on the optical axis of the injector lens, which resulted in a loss of power. All the laser diameter at 1/e² (D_L_) and laser power density P were estimated from average intensity profiles extracted with Fiji/ImageJ (Supplementary Fig. S10 and S11) and are expressed as the mean ± standard deviation.

For 1030-nm laser stimulation on the retina: D_L_ = 470 ± 70 µm. P = 0.56 ± 0.21 W.mm^-^², for 1030-nm photoacoustic stimulation with PDMS-CNT implants: D_L_ = 360 ± 60 µm. P = 0.39 ± 0.12 W.mm^-^², for 1030-nm photoacoustic stimulation with PDMS/CS/PDMS implants: D_L_ = 410 ± 45 µm. P = 0.29 ± 0.06 W.mm^-^².

The pupil of the eye of interest was dilated with a tropicamide-based eye drop solution (Mydriaticum®, Théa, France) before the first recording. Body temperature was monitored with a rectal probe and maintained using a heating blanket. Respiratory and heart rates were continuously monitored (TCMT, Minerve, France). After local application of lidocaine (4 mg.kg^-1^, Laocaïne®, MSD, France), the thinned skull was exposed and covered with ultrasound gel. The rats were scanned with a system dedicated to small animal ultrasound neuroimaging (Iconeus, Paris, France). Doppler vascular images were obtained using the Ultrafast Compound Doppler Imaging technique^47^. The probe was positioned coronally at Bregma - 6.5 mm in order to measure the cerebral blood volume (CBV) in the contralateral superior colliculus. Each frame was a compound plane wave frame^48^ resulting from the coherent summation of backscattered echoes obtained after successive tilted plane waves emissions. Then, the CBV signal was extracted from the tissue signal by filtering the image stacks with a dedicated spatiotemporal filter using Singular Value Decomposition^49^. Each transcranial Power Doppler image was obtained from 200 compounded frames acquired at 500 Hz frame rate.

#### Immunohistochemistry and image acquisition

After euthanasia, the eyes (n = 3 LE rats, n = 4 P23H rats) were enucleated and fixed with 4% paraformaldehyde (Cat no. 100496, Sigma-Aldrich) in 0.1 M phosphate buffer for 1 hour. The retinae were dissected and incubated in blocking solution (0.3% Triton X100 (Cat no. X100-500ML, Sigma-Aldrich), 10% Horse Serum (26050-070, Gibco) in phosphate buffer) with the primary antibodies (Iba1 (AB48004, Abcam), GFAP (Z0334, Dako), rhodopsin (MAB5316, Merck Millipore), cone arrestin (AB15282, Merck Millipore) for three days at room temperature. The secondary antibodies used to detect immunolabeling were goat anti-IgG conjugated to Alexa-Fluor 488, 594 or 647 (A212026, A21203, and A21447, Invitrogen).

The retinae were flat-mounted, and imaged using an Olympus FV3000 confocal microscope with a 20X objective (UPLXAPO20X NA 0.8 WD 0.6, Evident).

### Analysis of *in vivo* experiments

#### Analysis of OCT images

Mean retinal thickness next to and above the implant were measured with ImageJ on OCT images (diametral slices). The number of rats imaged at 30 dpi and later was lower than the number imaged at 7 dpi and 15 dpi because rats were used for terminal retinal stimulation recordings starting at 23 dpi.

#### Analysis of functional ultrasound imaging recordings

The correlation map of the CBV variations and the laser sequence for stimulation was computed by the manufacturer’s proprietary IcoStudio software (v1.5.2). A delay of either 2 or 3 s was computed in the calculation of the correlation to account for vascular delay (the chosen value maximizes the correlation). In correlation map displays (Fig. 7c), only significant pixels with a correlation threshold greater than 0.2 are shown. Maps with a correlation threshold of 0.1 are shown in Supplementary Fig. S12. A 29-by-29-pixel (29 pixels = 302 ± 10 µm) region of interest (ROI) was defined for each animal, centered on the peak intensity of the correlation map for full-field-white-light stimulation. Relative CBV variations (rCBV) were extracted in this ROI for all stimulation types. For each recording (15 laser stimulations), the cerebral blood flow (CBV) was normalized into a relative steady-state value (rCBV) and calculated as the following: rCBV = (CBV(t) - CBV_0_)/CBV_0_, where CBV(t) is the power doppler value t seconds after the start of laser sequence, and CBV_0_ is the baseline in the ROI. The baseline was defined as the mean power doppler value 5 seconds before the start of the laser sequence. The data was bootstrapped to calculate confidence intervals.

### Statistical analysis

Values are expressed as mean values ± standard error of the mean (SE) in figures and in the text, unless specified otherwise. Similarly, in scatter plots with error bars (Fig. 1, Fig. 4), data points and error bars represent the mean and the standard error of the mean, respectively.

Data normality was checked with SciPy normaltest. Data did not follow a normal distribution. Statistical significance was analyzed with Wilcoxon signed-rank tests and Mann-Whitney U tests (Fig. 2 to 6). Pearson correlation (Fig. 2 to 4) was computed to quantify the strength and direction of the linear relationship between two continuous variables. One-way ANOVA was used to test the effect of a single factor on the mean of a dependent variable (Fig. 6). Finally, to estimate confidence intervals for a statistic of unknown distribution (average rCBV variations in Fig. 7) we used bootstrapped estimation (1000 samples, 95% confidence intervals). Statistical tests are provided in the figure legends.

## Acknowledgements

This work was supported by the National Institute of Health, United States, through the BRAIN Initiative (BRAIN/NEI R21 EY035437 and NEI 1R21EY036579) to CY, by the Foundation Fighting Blindness (PPA-0922-0840-INSERM) to SP, by IHU FOReSIGHT, France, (ANR-18-IAHU-01) to SP, by the European Research Council (Grant Agreement 101045289) to SP, and by Axorus SAS, France, to SP and CY.

## Competing interests

This study was funded in part by the company Axorus SAS. J-DL and HM are major stakeholders in Axorus. CY and J-XC are minor stakeholders in Axorus.

## Author Contributions

## Equal contributions

A.L., Y.L. and TR.R, and S.P. and H.M.

A.L., Y.L., TR.R. and H.M. designed the experiments.

A.L., Y.L., TR.R., H.M., C.B., J.V., A.F., C-A.C., G.C., Y.Y., C.J., M.C. and J.D. carried out the experiments and analyzed the data.

J-D.L. and G.T. provided support for the experiments, study design and data analysis.

H.M., J-D.L., J-X.C., C.Y. and S.P. conceived the project, and contributed to study design and data interpretation.

A.L., Y.L., TR.R., C.Y.,H.M., and S.P. wrote the manuscript with input from all others.

## Data availability

Data are available from the corresponding author upon reasonable request.

## Code availability

The custom Python codes are available from the corresponding author upon request.

## Notes

### Summary of Updates

Modifications were made after a first round of peer review. Notably, we have performed additional experiments in which we 1. performed immunohistochemistry in implanted LE and P23H rats, in order to better assess the biocompatibility of our PA implant, and to evaluate the extent of photoreceptor degeneration in LE rats induced by the subretinal implantation; 2. applied L-AP4 and ACET pharmacological blockers to better assess the involvement of photoreceptors in retinal mechanosensitivity. The new experiments performed with pharmacological blockers led us to include a new subsection in the Results section, entitled "Mechanistic exploration of retinal mechanosensitivity". This section encompasses the new data with L-AP4 and ACET, as well as the data with (rs)-CPP and CNQX (previous Figure 2i). This data is presented in a new Figure 3, and all of the subsequent figures have been re-numbered appropriately. Moreover, following the most recent meeting of fUSI-BIDS (functional UltraSound Imaging - Brain Imaging Data Structure) users, the fUSI-BIDS community has chosen to adopt the acronym "fUSI" instead of "fUS" to describe functional ultrasound imaging, in order to avoid possible confusion with the "FUS" acronym, which stands for "Focused UltraSound". We have therefore changed all acronyms of "fUS" into "fUSI", in order to comply with those new guidelines.

