## Supplementary Information for "A flexible photoacoustic retinal prosthesis"

### A flexible high-precision photoacoustic retinal prosthesis

#### 1 Properties of the photoacoustic film

##### 1.1 Young's Modulus of PDMS/CS/PDMS photoacoustic film

We evaluated the Young's Modulus $E$ of the PDMS/CS/PDMS film using a tensile test (Supplementary Fig. S1) and the following equation:

$$E = \frac{Stress}{Strain} = \frac{FL}{A\Delta L},$$

where $F$ is the exerted force (N), $L$ is the original length (m), $A$ is the cross-sectional area ($m^2$), and $\Delta L$ is the change in the length (m).

Following our measurements, we calculated the PDMS film's Young's modulus to be 2.12 ± 0.10 MPa. It is two orders of magnitude lower than silicon-based implants (200-300 GPa), but remains several orders of magnitude higher than the retina's Young's modulus, which ranges between 0.5 kPa[1] and 25 kPa[2]. Therefore, our PA film is expected to provide better biocompatibility and induce less immune response than current implants.

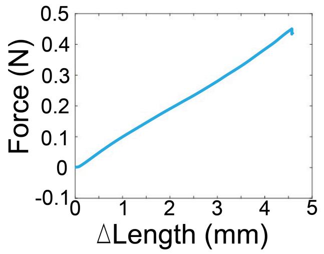


**Supplementary Figure S1. Raw data from the tensile test used to calculate Young's Modulus.** A rectangular piece of PDMS/CS/PDMS film was subjected to stretching, and deformation and force were measured.

##### 1.2 Peak pressure and energy conversion efficiency of the PDMS/CS/PDMS film

Pressure measurements were performed with a hydrophone (HGL-0085, Onda Corporation, USA), following the illumination of the photoacoustic (PA) film with a 1030-nm laser delivering 8-ns pulses with an energy of 7 µJ per pulse. A peak pressure of 146.2 kPa was measured at 900-µm away from the PDMS/CS/PDMS film, resulting in a photoacoustic conversion efficiency of 21 kPa.$\mu J^{-1}$ at 900-µm away from the PDMS/CS/PDMS film.

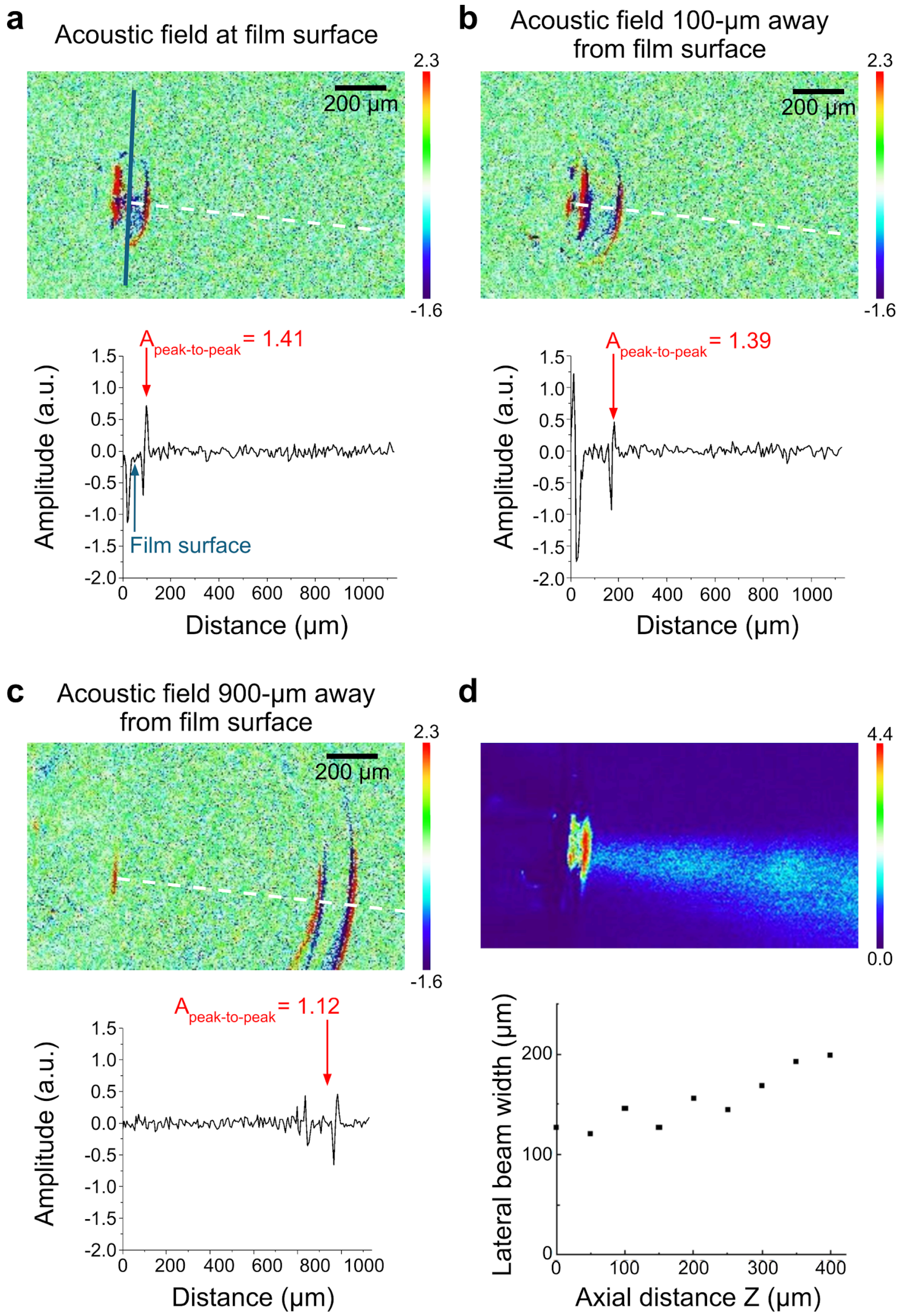


**Supplementary Figure S2. Ultrasound field generated by delivering laser pulses with a 200-µm optical fiber. (a)** Top: Acoustic wavefront at the surface of the film (blue line). The dotted white line shows the acoustic propagation direction. Bottom: acoustic amplitude as a function of distance (µm) from the candle soot (CS) layer. The film surface is 60-µm away from the CS layer. **(b)** Same as (a), but the acoustic wavefront is 100-µm away from the surface of the film. **(c)** Same as (a), but the acoustic wavefront is 900-µm away from the surface of the film. **(d)** Top: Acoustic field. Bottom: lateral beam width as a function of axial distance from the film surface. Lateral beam width ~150 µm at Z = 200 µm. As a reference, the average retinal thickness is 174.0 ± 2.3 µm in Long Evans rats and 72.3 ± 2.3 µm in P23H rats (Fig. 5c-d).

In order to estimate the photoacoustic conversion efficiency at the surface of the film, we mapped the PA field generated with a 200-µm fiber. The PA signal generated at the surface of the film had an amplitude of 1.41. At 100- and 900-µm away, the amplitude of the PA signal decayed to 1.39 and 1.12, respectively. Therefore, the photoacoustic conversion efficiency at the surface of the film is estimated to be 26 kPa.µJ$^{-1}$.

The energy conversion efficiency $E_{CE}$ is calculated using:

$$E_{CE} = E_A / E_0 ,$$

where $E_0$ is the optical energy (energy per pulse, $E_0$ = 7 µJ), and $E_A$ is the acoustic energy given by:

$$E_A = \frac{A}{\rho c}\int_0^{\infty} p^2(t)dt ,$$

given a laser spot area $A$ of 200-µm diameter (m$^2$), the density of water ρ = 998 kg.m$^{-3}$, the speed of sound in water $c$ = 1480 m.s$^{-1}$, and the peak-to-peak pressure of the acoustic wave (Pa).

Using our estimated photoacoustic conversion efficiency of 26 kPa.µJ$^{-1}$ at the film's surface and optical energy, we find an energy conversion efficiency value of $E_{CE}$ = 5.1 ×10$^{-5}$ % when applying a surface laser energy of 22 mJ.cm$^{-2}$.

**1.3 Optical properties of the CS layer**

To investigate the possibility of light leakage following the passage of laser pulses through the PDMS/CS/PDMS film, we measured the transmitted power using a power meter positioned behind the film. The measurements indicated negligible power transmission. We expect absorption to occur mainly at the CS layer. In order to measure a transmittance value in the detectable range of our spectrophotometer (UV-1900i from Shimadzu), we deposited a 200-nm-thick layer of CS on glass using flame deposition (same protocol as for PDMS/CS/PDMS implant fabrication). We then calculated the theoretical transmittance of the 4-µm-thick CS layer. At 1030 nm, transmission was measured at $T_1$ = 9.0 % for a CS layer thickness of $L_1$ = 200 nm (Supplementary Fig. S3). According to Beer-Lambert law, the expected light transmission $T(L)$ at 1030 nm for a CS layer thickness of $L$ = 4 µm is:

$$T(L) = 10^{(L/L_1 * logT_1)} .$$

We therefore find $T(L) = 1.1 \times 10^{-19}$ %.

Adsorption of the CS in the PDMS may affect how compact the layer is, and increase transmittance compared to the theoretically expected values.

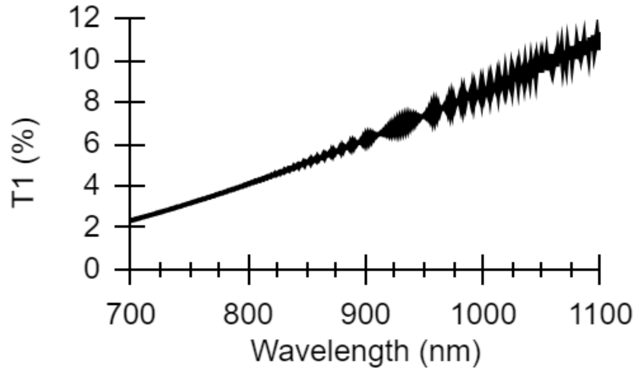


**Supplementary Figure S3. Transmittance ($T_1$) of a 200-nm-thick candle soot layer.**

##### 1.4 Characterization of PDMS-CNT film for *in vivo* photoacoustic retinal stimulation

We developed a second type of PA film for the *in vivo* experiments: a 40-µm thick PDMS-CNT film (Supplementary Fig. S4a). As for the PDMS/CS/PDMS film, photoacoustic signals were generated by delivering a pulsed 1030-nm laser and recorded with a hydrophone set 900-µm away from the film. The PDMS-CNT film emitted ultrasound with a peak pressure of 133 kPa for a laser energy of 10 µJ per pulse, resulting in photoacoustic conversion efficiency of 13.3 $kPa.\mu J^{-1}$ at 900-µm-away from the film surface. The PDMS-CNT film provides a central frequency of 10.9 MHz (compared to 42.2 MHz for the PDMS/CS/PDMS film) and -6 dB bandwidth of 5.9 to 15.8 MHz (Supplementary Fig. S4b).

Assuming the acoustic attenuation coefficient is 0.75 $dB.MHz^{-1}.cm^{-1}$ in the eye[3], the attenuation is estimated to be 1.09 fold at 900-µm away from the emitter. Therefore, the photoacoustic conversion efficiency at the surface of the film is estimated to be 14.4 $kPa.\mu J^{-1}$, and the energy conversion efficiency is $E_{CE} = 1.8 \times 10^{-4}$ % for a surface energy of 32 $mJ.cm^{-2}$.

In rats, the *in vivo* distance between the inner retina and the PA implant should be below 100 µm[4]. In the inner retina, the peak pressures of the acoustic waves generated with the PDMS-CNT and PDMS/CS/PDMS implants may therefore differ up to a factor 2.5 (66 $kPa.\mu J^{-1}$ vs 26 $kPa.\mu J^{-1}$). Still, we found significant and similar superior colliculus activation during PA stimulation with both implants (Fig. 6). The lack of measured difference between stimulation with both implants could be due to limitations of the recording method or saturation of superior colliculus responses due to strong stimulation.

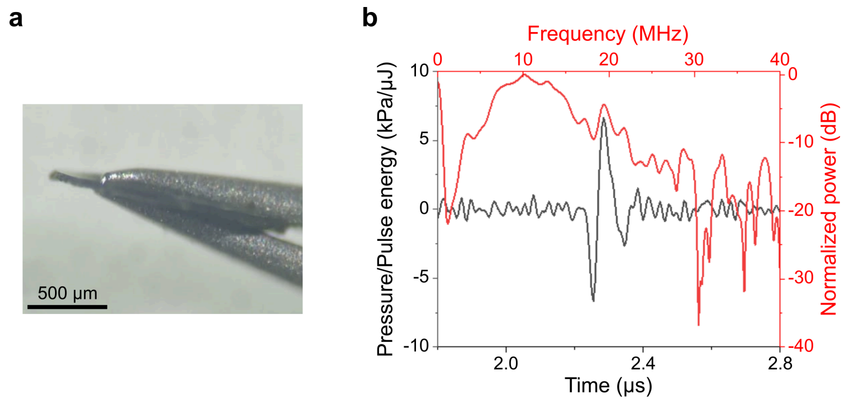


**Supplementary Figure S4. Characterization of the PDMS-CNT photoacoustic film. (a)** CNT-embedded PDMS with a thickness of 40 μm. **(b)** PA performance in the temporal domain (black) and frequency domain (red) of the photoacoustic films corresponding to films shown in (a).

#### 2 Safety considerations

##### 2.1 Mechanical index and spatial peak temporal average intensity

FDA safety regulations for ultrasound systems and transducers[5] for ophthalmic uses prescribe a mechanical index (*MI*) below 0.23, and a spatial peak temporal average intensity ($I_{SPTA}$) below 50 $mW.cm^{-2}$. These are defined as follows:

- $MI = NPP / \sqrt{f}$, where *NPP* is the negative peak pulse (acoustic) pressure (MPa) and *f* is the acoustic frequency (MHz);

- $I_{SPTA} = I * f_{rep}$, where $f_{rep}$ is the laser repetition frequency (Hz), and *I* is the pulse intensity integral ($W.cm^{-2}$);

- $I = \int_0^T p^2(t)dt / \rho c$, where ρ is the density of water ($kg.m^{-3}$), *c* is the speed of sound ($m.s^{-1}$), and *p* is the peak-to-peak pressure of the acoustic wave (Pa).

First, we calculated the upper bound values for *MI* and $I_{SPTA}$ based on the laser parameters that produce the strongest PA stimulation, i.e. those used for *ex vivo* photoacoustic stimulation: laser energy *E* = 10 µJ per pulse, delivered by a 200-µm-diameter fiber, with a repetition frequency of 3.5 kHz. *NPP* and peak-to-peak pressure values at the surface of the film are estimated from the experimentally measured photoacoustic conversion efficiency, as per Section 1.2.

Note that to establish these upper bound values, we assume that the laser spot diameter on the film during photoacoustic retinal stimulation is identical to that used for establishing the photoacoustic conversion efficiency of the PA film ($d_0 \simeq 200$ µm). In practice, the laser spot in *ex vivo* studies was closer to $d_1 = 300$ µm in diameter (mean laser energy density *P* = 0.52 $W.mm^{-2}$). At constant laser energy per pulse, both *NPP* and *p* are expected to decrease as laser spot diameter increases. For a rough estimate, we could consider that *NPP* and *p* values are linearly correlated to laser energy density, and so are to be divided by $(d_1 / d_0)^2 = 2.25$ to approximate experimental values.

For both PA films, the calculated upper-bound *MI* and $I_{SPTA}$ values are below FDA thresholds (Supplementary Table T1, green lines). The *MI* and $I_{SPTA}$ values for the PDMS/CS/PDMS film are 3- and 17-fold lower, respectively, than that of the PDMS-CNT film, making the former a safer option.

Second, we estimated lower bound values for *MI* and $I_{SPTA}$ (Supplementary Table T1, gray lines). *In vivo*, the mean laser spot size was 410 µm for PA stimulation with the PDMS/CS/PDMS film (mean laser energy density *P* = 0.39 $W.mm^{-2}$) and 360 µm (*P* = 0.29 $W.mm^{-2}$) for stimulation with the

PDMS-CNT film. Taking the laser parameters for *in vivo* stimulation, and assuming that *NPP* and *p* values are linearly correlated with laser energy density, we calculated lower bound values for *MI* and $I_{SPTA}$, which are 4- to 6-fold lower than the upper bound values.

| PA film | Laser spot diameter (µm) | *f* (MHz) | *NPP* (MPa) | ***MI*** | *I* (mW.cm$^{-2}$) | **$I_{SPTA}$ (mW.cm$^{-2}$)** |
|---|---|---|---|---|---|---|
| **PDMS/CS/ PDMS** | 200 (*ex vivo*) | 42.2 | 0.174 | **0.06** | 1.61E-05 | **0.056** |
| | 410 (*in vivo*) | 42.2 | 0.029 | **0.01** | 1.88E-06 | **0.011** |
| **PDMS-CNT** | 200 (*ex vivo*) | 10.9 | 0.330 | **0.10** | 2.68E-04 | **0.939** |
| | 360 (*in vivo*) | 10.9 | 0.071 | **0.02** | 4.10E-05 | **0.250** |

**Supplementary Table T1. Ultrasound characterization of photoacoustic films.** *NPP*: negative peak pressure, *MI*: mechanical index, *I*: ultrasound intensity, $I_{SPTA}$: spatial peak temporal average intensity. Green lines: upper bound values calculated using the laser parameters for *ex vivo* photoacoustic retinal stimulation and assuming a 200-µm laser spot diameter. Gray lines: lower bound values calculated using the laser parameters for *in vivo* photoacoustic retinal stimulation. The listed values comply with FDA thresholds[5] for *MI* and $I_{SPTA}$: $MI < 0.23$ and $I_{SPTA} < 50$ mW.cm$^{-2}$.

**2.2 Other safety concerns**

In the retina and, more generally, the eye, plasma formation due to high-energy laser pulses is a concern. In this study, the peak laser power, defined as surface pulse energy divided by pulse duration, is $P_{peak} \sim 10^5$ W.mm$^{-2}$. This value is three orders of magnitude below the threshold for plasma formation on the cornea, lens, and retina ($P_{peak} \sim 10^8$ W.mm$^{-2}$ for 6-ns laser pulses[6]).

### 3 Temperature increase

#### 3.1 Thermocouple measurement setup

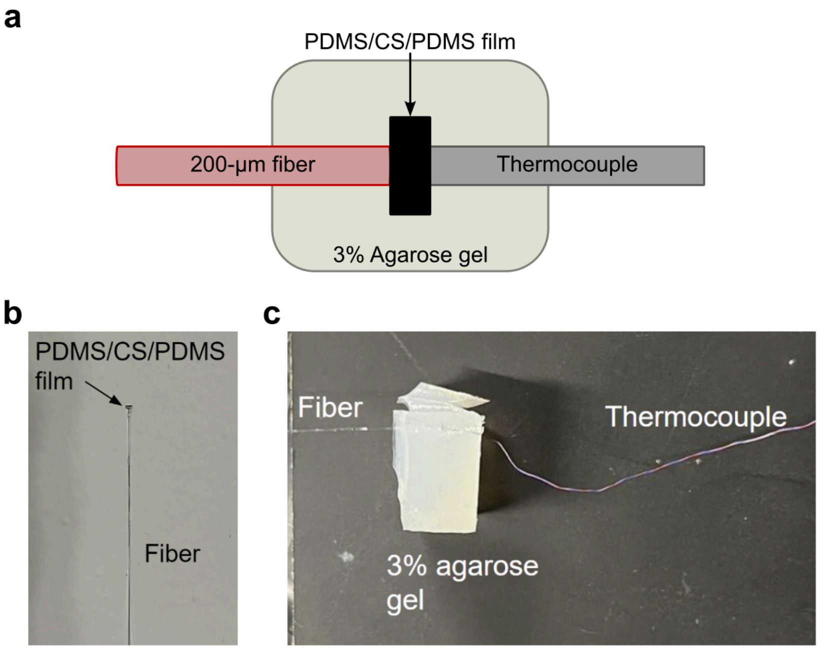


**Supplementary Figure S5. Thermocouple measurement setup. (a)** Schematic of the setup. **(b)** Photo of small PDMS/CS/PDMS film placed on the tip of a 200-µm optical fiber to facilitate alignment with the 200-µm thermocouple. **(c)** Photo of aligned fiber, PA film, and thermocouple sensor in 3% agarose gel.

#### 3.2 Transient temperature events with fast laser repetition rates

FDA safety guidelines for ultrasound systems and transducers[5] used in ophthalmic devices set the maximum local temperature increase to 1°C. Temperature increases with the laser stimulation parameters used *ex vivo* (Fig. 1g) and *in vivo* (Supplementary Fig. S6) have been measured to be below 1°C at the film surface using a thermocouple.

Transient temperature events in the 0.1-1 ms range could theoretically activate heat-sensitive TRPV1 and TRPV2 channels[7], which have activation thresholds of 43 °C[7–9] and 52 °C[7,9,10], respectively. However, the thermocouple's acquisition frequency of 2 kHz is not sufficient to capture those. These transient peaks would have to be induced by the individual laser pulses, which is incompatible with the absorber-to-cell distances in our system. Indeed, the transient component of the temperature rise, induced by laser pulses with a repetition frequency $f_{rep}$, propagates over a distance driven by the thermal diffusion length $\mu = \sqrt{(D/f_{rep})}$, where $D$ is the thermal diffusivity of the medium. If we consider $D_{water}$ = 0.14 mm².s$^{-1}$ (diffusivity for both pure[11] and carbon-loaded PDMS is in the range of 0.1 - 0.2 mm².s$^{-1}$) and repetition frequencies of 1.9 kHz and 6.1 kHz (the minimum and maximum laser repetition frequencies used in this study), then the thermal diffusion length is $\mu$ = 8.6 µm and 4.8 µm, respectively. With the PDMS/CS/PDMS film, the minimum distance of cells to the CS layer is 50 µm. Therefore, we do not expect transient temperature events to activate heat-sensitive channels.

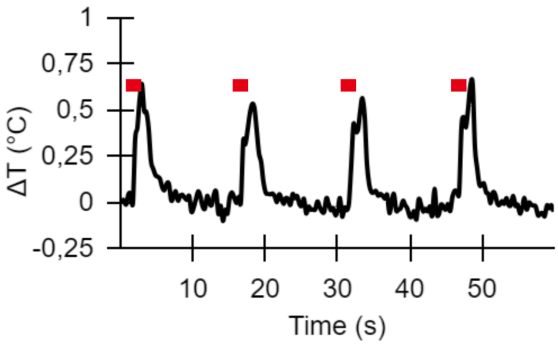


**Supplementary Figure S6. Temperature variation during *in vivo* stimulation conditions.** Temperature variation ($\Delta T$) at the film surface during 1030-nm laser irradiation. $P$ = 0.34 W.mm$^{-2}$. Red lines: laser ON. The same stimulus paradigm as for implant stimulation *in vivo*. Maximum temperature increase ($\Delta T$) of 0.64 °C.

**4 Comparison among retinal vision restoration technologies.**

| | This work | Argus II[12] | PRIMA by Pixium Vision[13] | POLYRETINA [14] | Ultrasound array[15] | GenSight's GS030[16] |
|---|---|---|---|---|---|---|
| **Working principle** | Photoacoustic | Electrical | Photovoltaic | Photovoltaic | Ultrasound | Optogenetics |
| **Physical spatial resolution** | 56-µm ultrasound field width | 200-µm pixel diameter and 575-µm pitch | 100-µm pixel size in patients, 20-µm in rodents[20] | 100-µm pixel diameter and 120-µm pitch | 81-µm ultrasound field width | Not applicable |
| **Field of view** | To be tested | 11° × 19° | 7° | 46.3° [17] | Theoretically > 40° | 8.2° |
| **Pixel density** | To be tested, theoretically > 2500 pixel/mm$^2$ | 3.6 pixel/mm$^2$ | 94.5 pixel/mm$^2$ in patients | 79.1 pixel/mm$^2$ [18] | 1.8 pixel/mm$^2$ on device | Not applicable |
| **Restored vision acuity** | To be tested | 20/1588[19] | 20/500 in patients, 20/100 in rodents[20] | To be tested, theoretically ~<20/400 based on resolution | To be tested | 20/249 |
| **Stimulation intensity** | 290 mW/mm | 1 mC/cm$^2$ | 3.5 mW/mm$^2$ | 33 µW/mm$^2$ | 3 MPa @ 4.4 MHz* | 13 mW/cm$^2$ |
| **Implant position** | Subretinal space | Epiretinal space | Subretinal space | Epiretinal space | Noninvasive | Intravitreal injection |
| **Development stage** | Preclinical in rodents | FDA approved | Multicentric clinical trial | Preclinical in minipigs | Preclinical in rodents | Pilot clinical trial |

* Mechanical index exceeds FDA safety requirement in the frequency range <14 MHz and ~18 MHz.

**Supplementary Table T2 A comparative table between our approach and other vision restoration technologies across multiple parameters.**

**5 Setup for photoacoustic field mapping**

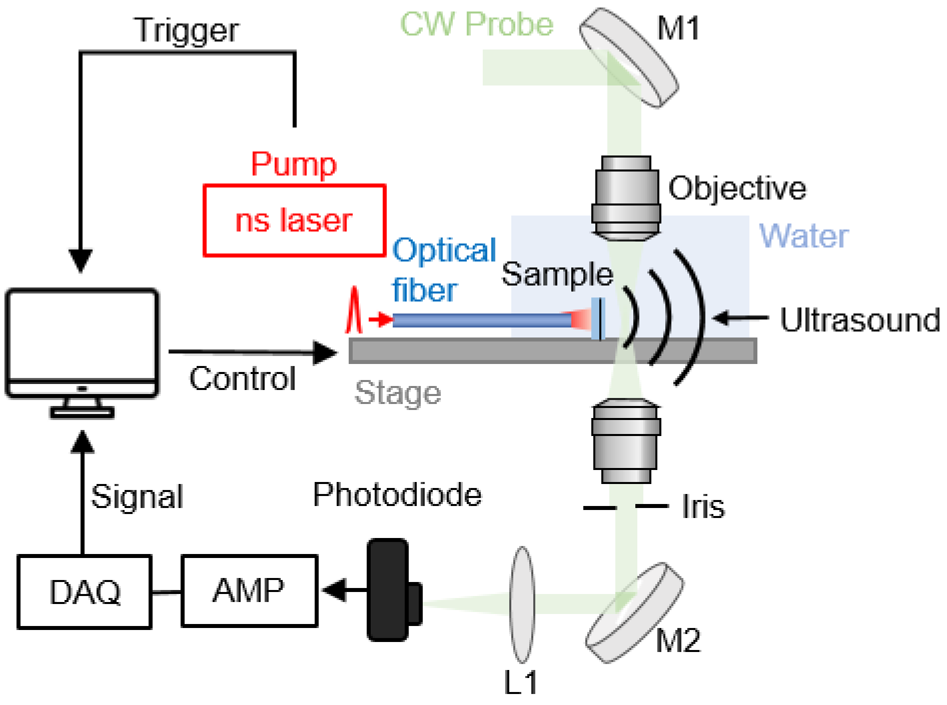


**Supplementary Figure S7. Setup for spatial-offset pump-probe imaging used to acquire the photoacoustic field**. Adapted with permission from Chen et al.[22]. The PDMS/CS/PDMS film was cut into small pieces and attached to the end of the optical fiber for characterization. The nanosecond pump laser, delivered through an optical fiber, generated the photoacoustic signal. An orthogonally aligned probe laser detected the resulting ultrasound field by sensing changes in optical signal amplitude caused by ultrasound-induced density variations in the medium. More details about the setup can be found in Chen et al.[22].

#### 6 Photoelectric effect of laser on MEA

In this study, we used a multi-electrode array (MEA) to measure retinal ganglion cell activity. We investigated whether the laser resulted in the electrical signals due to the photoelectric effect. First, we examined the effect of the laser transmitted through the PA film. The configuration of the measurement was the same as for *ex vivo* experiments, except with no retina (Supplementary Fig. S7a, top). Following the photoactivation of the PA film, a low frequency electrical signal was measured by the MEA at the onset of photoacoustic stimulation (Supplementary Fig. S7a, bottom). This signal is too slow to be photoelectric effect, and is possibly an indirect detection of induced temperature variations. When the 1030-nm pulsed laser directly illuminates the MEA (Supplementary Fig. S7b-d), it generates a strong photoelectric signal, with individual voltage peaks for each laser pulse (Supplementary Fig. S7d). At comparable energy density, the photoelectric signal is much stronger than the slow wave signal generated by the PA film. The lack of photoelectric effect when the MEA is covered by the PA film is coherent with the expected low light transmission of the film.

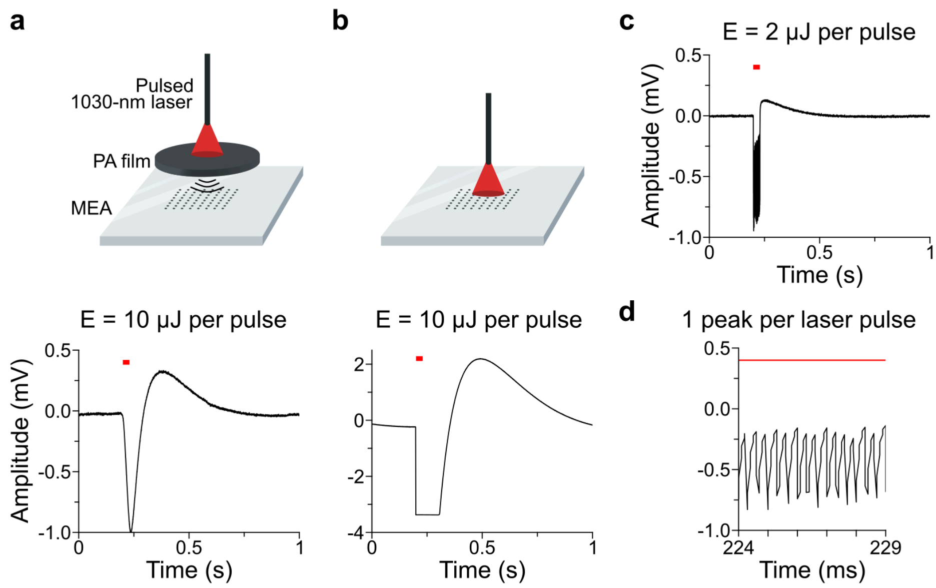


**Supplementary Figure S8. Raw voltage recording on the MEA electrode closest to the center of the laser beam with and without PDMS/CS/PDMS film. (a)** Top: setup with photoacoustic (PA) film between laser and multielectrode array (MEA). Bottom: voltage recording from the MEA electrode on which the laser is centered (greatest signal amplitude on MEA). Red horizontal line: laser on. Laser parameters: energy per pulse $E$ = 10 μJ, repetition rate $f_{rep}$ = 2.94 kHz, burst duration $d_b$ = 30 ms. These parameters are in the high end of parameters tested during ex vivo measurements. **(b)** Top:

setup with the laser directly illuminating the MEA. Bottom: same as (a). Same laser parameters as (a). Electrode saturation occurs for voltage signals greater than 3.4 µV. **(c)** Same setup as (b). Laser parameters: $E$ = 2 µJ per pulse, $d_b$ = 15 ms. $E$ divided by 5 and $d_b$ divided by 2 to avoid electrode saturation and obtain voltage signals of comparable amplitude to (a). **(d)** A magnified version of the X axis of (c). In 5 ms, the laser generates 14-15 optical pulses at 2.94 kHz. We can count 14 voltage peaks in the MEA recording.

#### 7 *Ex vivo* RGC count per stimulation site

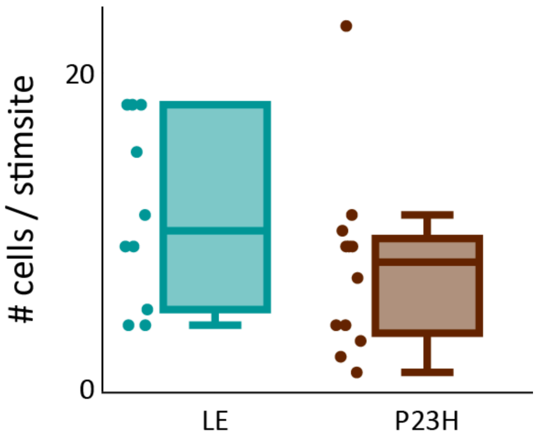


**Supplementary Figure S9. RGC cell count per stimulation site.** Number of RGCs with baseline activity in a 600-µm-diameter area (the "stimulation area") centered on the 300-µm-diameter area illuminated by the laser during stimulation. Each dot represents an individual stimulation area. LE: 10 stimulation areas from 4 retinas, n = 11 ± 5 cells per stimulation area (mean ± S.D.). P23H: 12 stimulation areas from 5 retinas, n = 7 ± 5 cells per stimulation area.

#### 8 *In vivo* - laser irradiance calculation

##### 8.1 Characterization of the laser beam exiting the laser injector

The laser injector from the MICRON 810-nm Image-Guided Laser modality is designed to project a laser spot of similar size to the diameter of the optical fiber used for delivery at the focal point of the injector lens. At the focal plane (7 mm from the injector lens, equivalent to the average diameter of a rat's eye), the measured laser beam radius, delivered with a 200-µm-diameter fiber, was $w_1$ = 162 µm.

A continuous laser with a repetition frequency $f_{rep}$ = 6.1 kHz was applied to the laser injector. The power exiting the laser injector $P_0$ was measured with a power meter. The resulting energy per pulse $E_{p0}$ was calculated using $E_{p0} = P_0 / f_{rep}$ = 15 µJ/pulse.

When the laser beam exits the injector along the optical axis ("on-axis" laser beam), e.g. through the center of the injector's lens, the power is concentrated at the laser focal spot. When the laser beam exits off the axis ("off-axis" laser beam, with $r$ the distance to the optical axis), the laser focal spot holds only a fraction of the total laser power. The ratio $R_P(r) = I(r) / I(0)$, representing the laser intensity at the focal spot for on- and off-axis configurations, was measured from images of the laser spot exiting the injector (Supplementary. Fig. S9a) using Image J by extracting the integral pixel value of the laser profile (cutoff at $1/e^2$ of maximum). The resulting off-axis energy per pulse can then be calculated with: $E_p(r) = R_P(r) * P_0$.

For a given laser spot, the beam radius at $1/e^2$ is calculated using Gaussian interpolation of the laser intensity profile (Supplementary. Fig. S9b). In the on-axis configuration ($r$ = 0), laser radius is $w_1$ = 162 µm. Resulting power density is $P_1 = f_{rep} * E_{p1}/(\pi w_1^2)$, with $E_{p1}$ = 15 µJ per pulse. $P_1$ = 1.11 W.mm$^{-2}$. In the off-axis configuration ($r$ = 292 µm), $w_2$ = 120 µm. The measured intensity ratio is $R_p(r)$ = 0.28 (Supplementary Fig. S9c). As a result, for $r$ = 292 µm, energy per pulse at focal point is $E_{p2} = R_p(r) * E_{p1}$ = 4.2 µJ and power density is $P_2 = E_{p2} / f_{rep}$ = 0.57 W.mm$^{-2}$.

In this example, the power density is approximately halved in the off-center position.

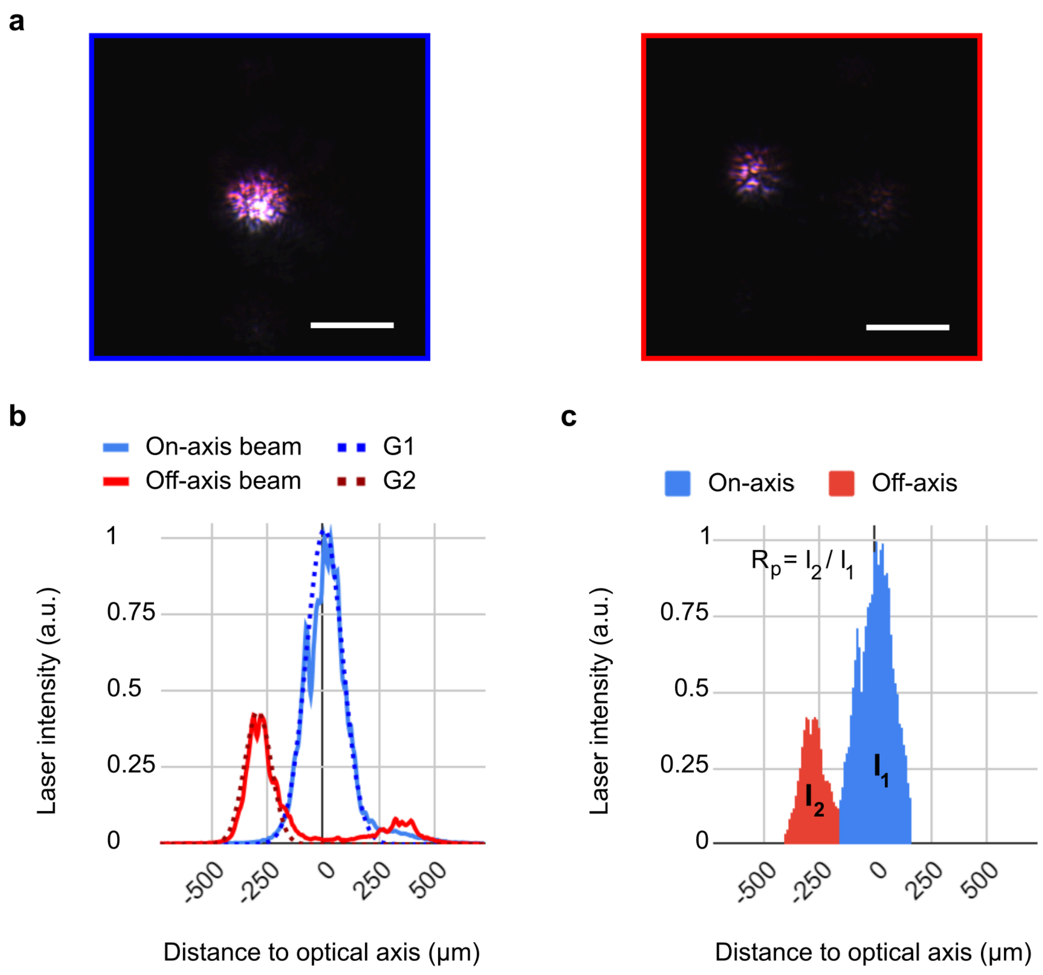


**Supplementary Figure S10. Estimation of laser power density exiting the laser injector. (a)** Images of the 1030-nm laser beam exiting the injector. Left: on-axis beam, $r = 0$. Right: off-axis beam, $r = 292$ µm. Scale bar: 500 µm. **(b)** Laser intensity profiles at the focal spot of an on-axis and off-axis 1030-nm laser beam. Experimental profile (continuous line) and Gaussian interpolations (dotted lines). **(c)** Integrals $I_1$ and $I_2$ (cutoff at $1/e^2$ of maximum) of the experimental laser profiles, respectively on-axis and off-axis. In this specific example, $R_p = 0.28$.

#### 8.2 Characterization of the laser beam on the retina and on the implant

In the previous section, we estimated the laser power density at the focal point when the laser beam exits the laser injector off-axis. In practice, the rat retina is not in the focal plane of the injector lens during *in vivo* stimulations. When the PA implant or the rat retina is closer to the injector lens than the focal planes, the diameter *D* of the laser beam on the implant or retina will be larger than the laser spot diameter at the focal plane, whether the beam is off- or on-axis (Supplementary Fig. S10a). This will further reduce the laser power density.

When using a laser at repetition rate $f_{rep}$ = 6.1 kHz and an energy per pulse $E_{p1}$ = 15 µJ/pulse, the resulting laser power density at the focal plane (in an on-axis configuration) is $P_1 = f_{rep} * E_{p1}/(\pi * w_1^2)$ = 1 W.mm$^{-2}$, with $w_1$ = 162 µm as described in the previous section.

The assumptions made to calculate laser power density (W.mm$^{-2}$) during stimulation on the retina or on the implant were the following:

- for $E_P(r)$:
  - no light absorption in the eye (see section 6.3);
  - $E_p$ = 15 µJ/ per pulse when.when the laser beam is on-axis;
  - no reflection of laser light on the implant. All the injected light is considered absorbed by the implant and converted into acoustic or thermal energy;
- The size of the imaged laser spot (on camera) is equal to the size of the spot on the PA implant.

In addition to optical aberrations due to the injector lens, there may also be spherical aberrations due to the biological lens[22].

We experimentally measured a laser beam radius of $w_{eye}$ = 195 µm from eye fundus images with the laser on (Fig 6b(ii), Supplementary Fig. S10b). This suggests that the retina is in front of the injector lens' focal point. When using the same laser repetition rate ($f_{rep}$ = 6.1 kHz) and energy per pulse ($E_{p1}$ = 15 µJ per pulse) as the previous paragraph, the resulting power density is $P_{eye}$ = 0.69 W.mm$^{-2}$.

Control experiments with a 595-nm and a 1030-nm laser used to directly stimulate the retina were all done in an on-axis configuration. Experiments on implants had to be performed in off-axis configurations to align the laser on the 1-mm-diameter implant.

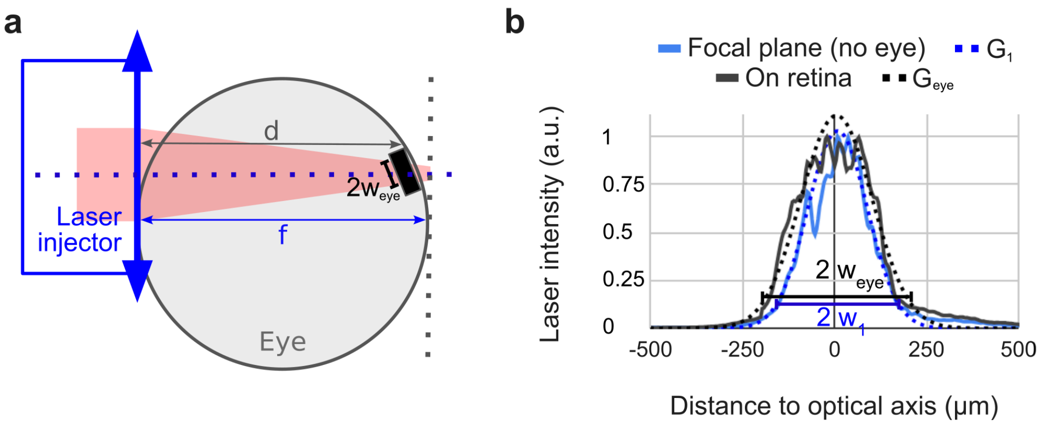


**Supplementary Figure S11. Estimation of the laser density on the retina or implant. (a)** Schematics of an injected laser beam where the injector lens focal plane is behind the implant. $f$ = 7 mm, $d < f$.

Laser spot diameter on implant *D = 2*$w_{eye}$. **(b)** Laser intensity profiles (solid line) obtained from eye fundus, and gaussian fits (dotted line) of on-axis laser beam at focal plane (blue) and on a rat retina (black). Laser diameter *D = 2w*, with laser beam radius a $1/e^2$.

**8.3 Neglect of laser absorption in the eye**

Before reaching the implant, the 1030-nm laser light goes through the aqueous humor, the lens and the vitreous. In humans, more than 90% of 1030-nm light is transmitted through the aqueous humor, and more than 80 % is transmitted through the vitreous[22]. Given that the thicknesses of the aqueous humor and the vitreous are much smaller in rats than in humans, the total light absorption in a rat's eye is therefore low enough (~ 20%) that we chose to assume no light was absorbed by the rat eyeball.

#### 9 Superior colliculus activation following photoacoustic retinal stimulation of *in vivo* LE retinae.

Pixels with a correlation threshold increase greater than 0.2 between the different stimulations and the relative cerebral blood volume variations (rCBV) are displayed on the correlation map in Fig. 6c. For lower correlation values, the increase of rCBV compared to the baseline is not significant. Supplementary Fig. S11 shows additional correlation maps (bottom line) with a 0.1 minimum pixel correlation threshold. With this lowered threshold, pixels appear in the contralateral Superior Colliculus (cSC) of the brain for 1030-nm laser stimulation of the retina. This suggests that for higher laser energy levels, the cSC may significantly respond to infrared pulsed stimulation.

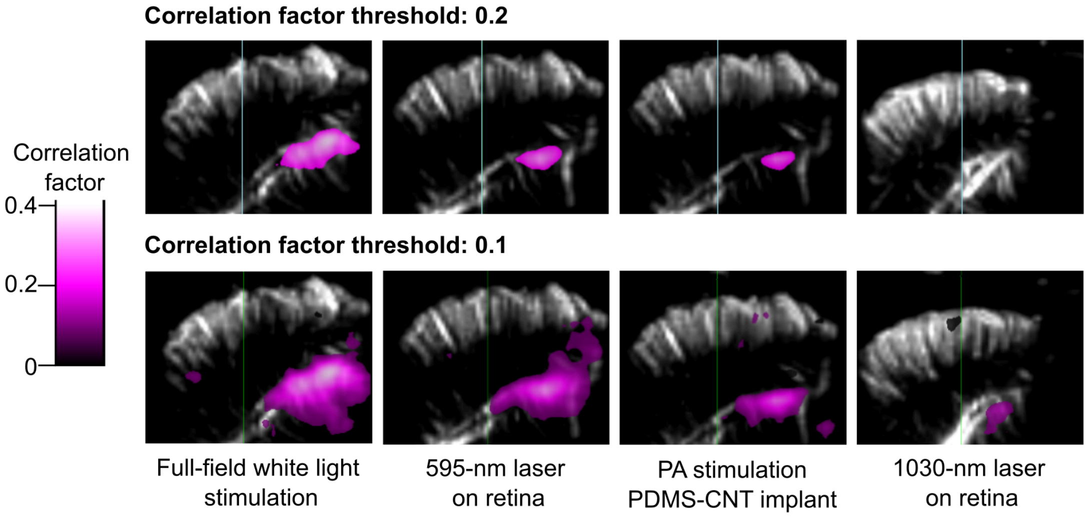


**Supplementary Figure S12. Superior colliculus activation following photoacoustic retinal stimulation of in vivo LE retinae.** Brain slice of one rat (coronal plane, left hemisphere, AP Bregma -6.5 mm) with correlation maps displaying cSC activation for a single recording (15 stimulations). Top row: correlation threshold between rCBV and the laser sequence is 0.2 (same as Fig. 6c). Bottom row: correlation threshold is 0.1.
